# GeneCover: A Combinatorial Approach for Label-free Marker Gene Selection

**DOI:** 10.1101/2024.10.30.621151

**Authors:** An Wang, Stephanie Hicks, Donald Geman, Laurent Younes

## Abstract

The selection of marker gene panels is critical for capturing the cellular and spatial heterogeneity in the expanding atlases of single-cell RNA sequencing (scRNA-seq) and spatial transcriptomics data. Most current approaches to marker gene selection operate in a label-based framework, which is inherently limited by its dependency on predefined cell type labels or clustering results. In contrast, existing label-free methods often struggle to identify genes that characterize rare cell types or subtle spatial patterns, and they frequently fail to scale efficiently with large datasets. Here, we introduce geneCover, a label-free combinatorial method that selects an optimal panel of minimally redundant marker genes based on gene-gene correlations. Our method demonstrates excellent scalability to large datasets and identifies marker gene panels that capture distinct correlation structures across the transcriptome. This allows geneCover to distinguish cell states in various tissues of living organisms effectively, including those associated with rare or otherwise difficult-to-identify cell types. We evaluate the performance of geneCover across various scRNA-seq and spatial transcriptomics datasets, comparing it to other label-free algorithms to highlight its utility and potential in diverse biological contexts.

## 1 Introduction

The identification of marker genes plays a critical role in advancing our understanding of cellular and spatial heterogeneity at the transcriptomic level. With the continuous expansion of scRNA-seq and spatial omics data, the ability to identify informative marker gene panels has become essential for characterizing distinct cell states and their spatial distribution within tissues. These insights are fundamental for unraveling complex biological processes and for constructing comprehensive cellular atlases across various tissues and organisms.

Current approaches to marker gene selection can be broadly categorized into three types: generative, label-based and label-free. Generative methods [1, 2, 3, 4, 5] build statistical models of gene expression in which cell types and marker genes enter as latent variables. They typically rely on heavy computation and are not easily scalable. Label-based methods rely on predefined cell type labels to identify marker genes that differentiate between these cell types. Notable examples of such label-based methods include Seurat [6] differentially expressed gene (DEG) analysis, scGenefit [7], RankCorr [8], and CellCover [9]. While effective, these methods have inherent limitations due to their reliance on clustering-based cell type labeling or manual annotation. Clustering-based labeling typically focuses on identifying cell sub-populations that exhibit significant variability at the transcriptomic level, as principal component analysis (PCA) is often applied prior to graph-based clustering. This emphasis on high-variability features may obscure the detection of subtle or rare cell types, as the genes characterizing these populations often do not display dominant variability patterns. Moreover, manual annotation at single-cell resolution requires expert knowledge, making the process both time-consuming and resource-intensive.

In contrast, most existing label-free marker gene selection methods, which do not rely on predefined cell type labels, adopt an imputation-based objective. These methods aim to select gene panels that effectively recover the underlying structure of the entire transcriptome. For example, PERSIST [10] selects genes that are maximally predictive of the overall gene expression profile using a concrete autoencoder network. Similarly, SCMER [11] identifies an optimal gene set by preserving the graph structure defined by pairwise cell similarity scores. GeneBasis [12] employs a greedy algorithm to select a gene panel that maintains the distance between the data manifold of the full transcriptome and that formed by the selected genes. Additionally, DUBStepR [13] uses stepwise regression on the gene-gene correlation matrix to predict the correlation matrix of the full gene set from the selected genes, iteratively regressing out the gene that explains the largest amount of variance in the residuals from the previous step. While these imputation-based methods provide effective unsupervised solutions by selecting features that preserve the global structure of complex, high-dimensional omics data, the gene panels they produce often reflect a broad, global representation of the data, yet are less sensitive to small cell populations. This is because the data structure these methods seek to recover is predominantly influenced by genes with high variability, or, as in the case of DUBStepR, the selection process itself is driven by explained variance. Consequently, these imputation-based approaches may overlook genes that are crucial for identifying rare cell types or capturing fine spatial organization, limiting their ability to detect nuanced biological signals.

Given the preceding discussion, both label-based methods and imputation-based label-free approaches face limitations in capturing biological signatures from rare sources of variability. To facilitate the discovery of genes associated with all sources of transcriptomic variability, we introduce geneCover, a label-free correlation-based marker gene selection method designed for single-cell RNA-seq and spatial transcriptomics data. GeneCover is motivated by the observation that, within highly heterogeneous gene expression profiles, groups of genes that characterize specific cell states or spatial organizations exhibit similar expression patterns, forming unique correlation structures among the transcriptome [13]. It highlights the potential of correlation-based methods for capturing both local and global signals by identifying distinct correlation groups formed by genes associated with rare and major cell types separately.

To capture these diverse genome-wide correlation structures, geneCover employs a minimal set-covering approach applied to the pairwise gene correlation matrix. This novel combinatorial strategy provides a globally optimal solution for the identification of minimally redundant gene panels that represent each distinct correlation structure, effectively characterizing unique spatial and cellular expression patterns at both local and global scales. By focusing on gene-gene correlations, geneCover is capable of identifying markers with subtle variations in transcriptomic activity that imputation-based methods often overlook, thereby providing a comprehensive tool for studying complex biological systems.

We demonstrate that geneCover enhances the detection of transcriptionally distinct cell types and spatially organized cell populations. To evaluate the performance of geneCover, we conducted an extensive comparison with leading imputation-based label-free marker gene selection methods across multiple scRNA-seq and spatial transcriptomics datasets. GeneCover outperforms or matches the best imputation-based methods in recovering cell identities and spatial organization. Notably, geneCover increases the resolution of rare cell types and subtle spatial structures compared to imputation-based approaches. As illustrations, we show that geneCover successfully distinguishes the hippocampal subfields in a mouse brain Visium HD dataset [14, 15], a level of resolution that other imputation-based methods fail to achieve. In a second example, analyzing a breast cancer scRNA-seq dataset [16], we show that geneCover identifies a transcriptionally distinct immune cell subpopulation, potentially linked to dendritic cells, which is absent from the original cell type annotation.

In addition to its improved resolution, geneCover scales more efficiently compared to existing label-free approaches, allowing it to handle large and complex datasets with significantly reduced computational overhead compared to competing methods. This scalability makes geneCover well-suited for use with the latest Visium HD, enabling efficient exploration of cellular heterogeneity at an unprecedented level of detail. Furthermore, geneCover enables marker gene selection across multiple tissue samples, allowing the identification of transcriptional programs shared across biological replicates. We demonstrate that geneCover can effectively uncover the conserved spatial organization of human cortical layers across multiple DLPFC donors, highlighting its potential to enhance cross-sample biological insights in spatial transcriptomics.

## 2 Methods

### Notation

In the analysis of spatial transcriptomics data, each tissue section is decomposed into a collection of discrete locations, referred to as “spots,” at different cellular resolutions depending on the sequencing technology. For scRNA-seq data, each individual cell serves as the fundamental unit of analysis. To unify these concepts, we denote both spots and cells as basic units in the target dataset, indexed from 1 to *N*. The gene expression levels across these units are represented as 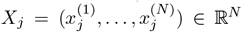, where *j* ∈ ⟦*d*⟧ = {1, …, *d*} denotes the *j*^*th*^ gene out of the total *d* genes, and 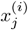 is the expression of gene *j* in the *i*^*th*^ unit. We can convert *X*_*j*_ to its rank representation and denote it as *R*(*X*_*j*_), where the *i*^th^ element of *R*(*X*_*j*_) records the rank of 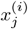 in *X*_*j*_.

Each gene *j* is also associated with a weight *w*_*j*_, reflecting the cost of its inclusion in the marker panel. To account for the gene-gene correlations, we define *ρ*(*j, j*′) as a measure of the correlation between *X*_*j*_ and *X*_*j*_′, where *j, j*′ ∈ ⟦*d*⟧. By default, we will set *w*_*j*_ = 1 for *j* ∈ ⟦*d*⟧ and use Spearman’s correlation

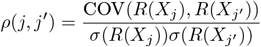

as the correlation measure. Given a subset of genes *G* ⊆ ⟦*d*⟧ ^1^ and a correlation threshold λ, we define the neighborhood of gene *j* ∈ *G* as 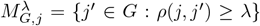. This correlation structure is encoded in a binary adjacency matrix 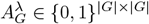 such that 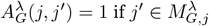 and 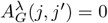 otherwise. Similarly, we denote ***ρ***_***G***_ and ***w***_***G***_ as the correlation matrix and weight vector with genes in *G*.

### Minimal Weight Set Covering

The minimal-weight set covering problem aims to identify a subset of genes *J* ⊆ *G* that covers the remaining transcriptome while minimizing the total weights of the selected genes. Let *M*_*G*,*j*_ denote the set of neighboring genes for each *j* ∈ *G*. A minimal set cover is a subset *J ⊂ G* of minimum cardinality such that ⋃_*j*∈ *J*_ *M*_*G*,*j*_ = *G*. In other words, the covering requires each gene *j*′ ∈ *G* to belong to at least one *M*_*G*,*j*_ where *j* ∈ *J*. The weighted version of the problem minimizes Σ_*j*∈ *J*_ *w*_*j*_ over all covering sets *J*.

Minimal weight set covering is a classical problem in combinatorial optimization [17] and can be formulated as an integer programming problem. Introduce the binary vector ***u*** ∈ {0, 1}^|*G*|^, where *u*_*j*_ = 1 indicates that gene *j* ∈ *G* is selected for the marker panel. Given the thresholding parameter λ, the integer programming formulation is

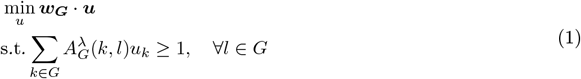

where the objective function minimizes the total weight of the selected genes while ensuring that each gene *l* ∈ *G* is covered—i.e., correlated above the threshold λ—by at least one of the selected marker genes. We solve this integer programming problem using the Gurobi optimizer [18]. The optimal covering set is *J*_*G*_(λ) = {*j* ∈ *G* : *u*_*j*_ = 1}.

### Refinement and Size Adjustment

The optimal solution *J*_*G*_(λ) obtained from the integer programming formulation includes genes that only cover themselves, i.e., 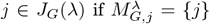. However, genes that correlate only with themselves or with few other genes often exhibit noisy expression patterns and contribute limited biological insight. To increase the robustness of our marker panel, we exclude genes *j* ∈ *J*_*G*_(λ) that cover fewer than *m* genes where *m >* 1. The final marker gene panel is thus refined as

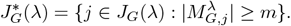

To obtain a marker gene panel 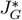 of pre-defined size *k*, we perform a binary search on parameter λ until 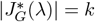.

### Expansion

One may also wish to expand the marker panel to capture a broader set of genes representing multiple genes from each correlated gene group. Since minimal set covering identifies a compact set of genes that characterize distinct correlated gene groups, we can expand the panel by iteratively selecting additional genes from these groups. To achieve this, at each iteration *t* + 1, we remove the lastest optimal marker panel 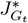 from the set of genes considered in the previous iteration *G*_*t*_ and then run the minimal set covering on the remaining genes 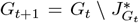 with marker panel refinement to identify a new panel 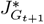 of pre-selected size that captures additional genes from the remaining correlated gene groups. Since each iteration is performed on the reduced gene set, this process may also uncover new correlated gene groups that were not prioritized in earlier iterations, thereby providing a more comprehensive representation of gene expression variability. The expanded marker panel is obtained by repeating this process over multiple iterations until the desired panel size or coverage is achieved.

### GeneCover Algorithm

The geneCover algorithm takes as input the whole-transcriptome gene-gene correlation matrix ***ρ*** ∈ ℝ^*d*×*d*^, a positive weight vector ***w*** ∈ ℝ^*d*^, a target subset of the transcriptome *G* ⊆ ⟦*d*⟧, the marker panel refinement parameter *m* and the pre-selected marker panel size *k* or a non-negative sequence of sizes {*k*_*t*_}_1:*T*_ for successive expansions. It is fully described in Algorithm 1.

### 3 Results

To systematically evaluate the performance of geneCover, we applied our method across a range of distinct biological systems captured by both scRNA-seq and spatial transcriptomics datasets from multiple protocols (See Supplementary Section 1 for details of dataset processing and marker panels generation from other label-free methods):

– The **DLPFC** dataset [19] offers spatial mapping of gene expression across the six layers of the human dorsolateral prefrontal cortex using the 10x Genomics Visium platform. With manual histological layer annotation, this dataset is commonly used as a benchmark for evaluating spatial transcriptomics methods.
– The **CBMC** CITE-Seq dataset [20] is a multimodal single-cell analysis derived from cord blood mononuclear cells (CBMCs). It combines cell-surface protein expression with transcriptomic data, offering a rich source of immune cell population information in the cord blood system. For our analysis, we focus exclusively on RNA expression data.

#### Algorithm 1

GeneCover

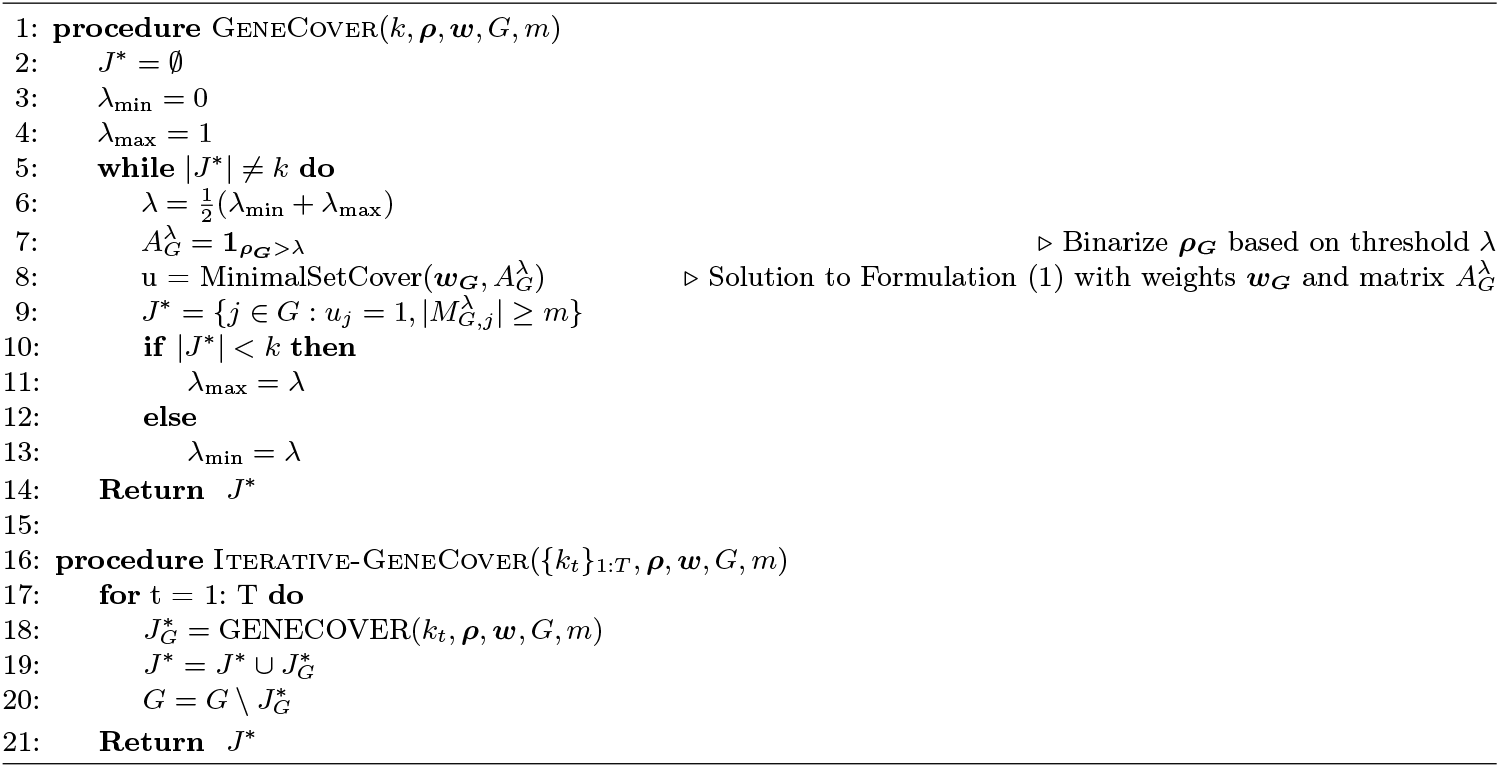

– The **mouse brain Visium HD** dataset [15] is derived from an FFPE brain tissue block of an eightweek-old male mouse, providing a high-resolution, whole-transcriptome spatial mapping of multiple brain regions.
– The **scFFPE breast cancer** dataset [16] provides a detailed molecular characterization of breast cancer, identifying 15 distinct cell types to improve understanding of tumor progression and immune interactions.

The following sections present an experimental evaluation of geneCover. First, we compare geneCover with other leading label-free marker gene selection methods on the DLPFC dataset, demonstrating its effectiveness in identifying spatially organized gene expression patterns. Next, we illustrate that geneCover enhances the resolution of single-cell and spatial transcriptomic discoveries, focusing on its ability to uncover nuanced cell types and spatial organizations in the CBMC, mouse brain Visium HD, and scFFPE breast cancer dataset. We then discuss the scalability of geneCover, showcasing its ability to efficiently handle large datasets. Following this, we analyze the robustness of geneCover’s hyperparameter selection, evaluating the stability of our clustering results across different hyperparameter settings. Lastly, we introduce a generalized geneCover framework that enables marker gene selection across multiple samples, highlighting its ability to identify conserved cellular programs.

### 3.1 Relative Performance in Recovering Cell Identities

To benchmark the performance of geneCover against other label-free gene selection methods, we conducted an experiment using the DLPFC dataset (sample #151673). We compared geneCover with five other methods: geneBasis, PERSIST, SCMER, and DUBStepR, which are label-free imputation-based marker gene selection methods, as well as Highly Variable Genes (HVGs). For each method, we obtained marker gene panels from the gene expression profile of the DLPFC dataset and restricted the log-normalized count matrix to the selected genes. We then performed principal component analysis (PCA), retaining 50 principal components based on these gene panels. To assess the clustering performance, we applied the Leiden [21] algorithm in SCANPY [22] with default parameters to these principal components across 30 random seeds and computed the average normalized mutual information (NMI) between the resulting clusters and the manually annotated histological layers. This benchmarking procedure allowed us to quantify how well each method’s marker gene panel could recover the morphological structure of the prefrontal cortex.

The results, shown in Figure 1A, indicate that geneCover consistently performs at similar or higher levels compared to other methods across different marker panel sizes, with geneBasis being the closest competitor. Specifically, geneCover maintains this performance advantage across different marker panel sizes, demonstrating its robustness and effectiveness in recovering spatially organized structures in the DLPFC.

**Fig. 1:**
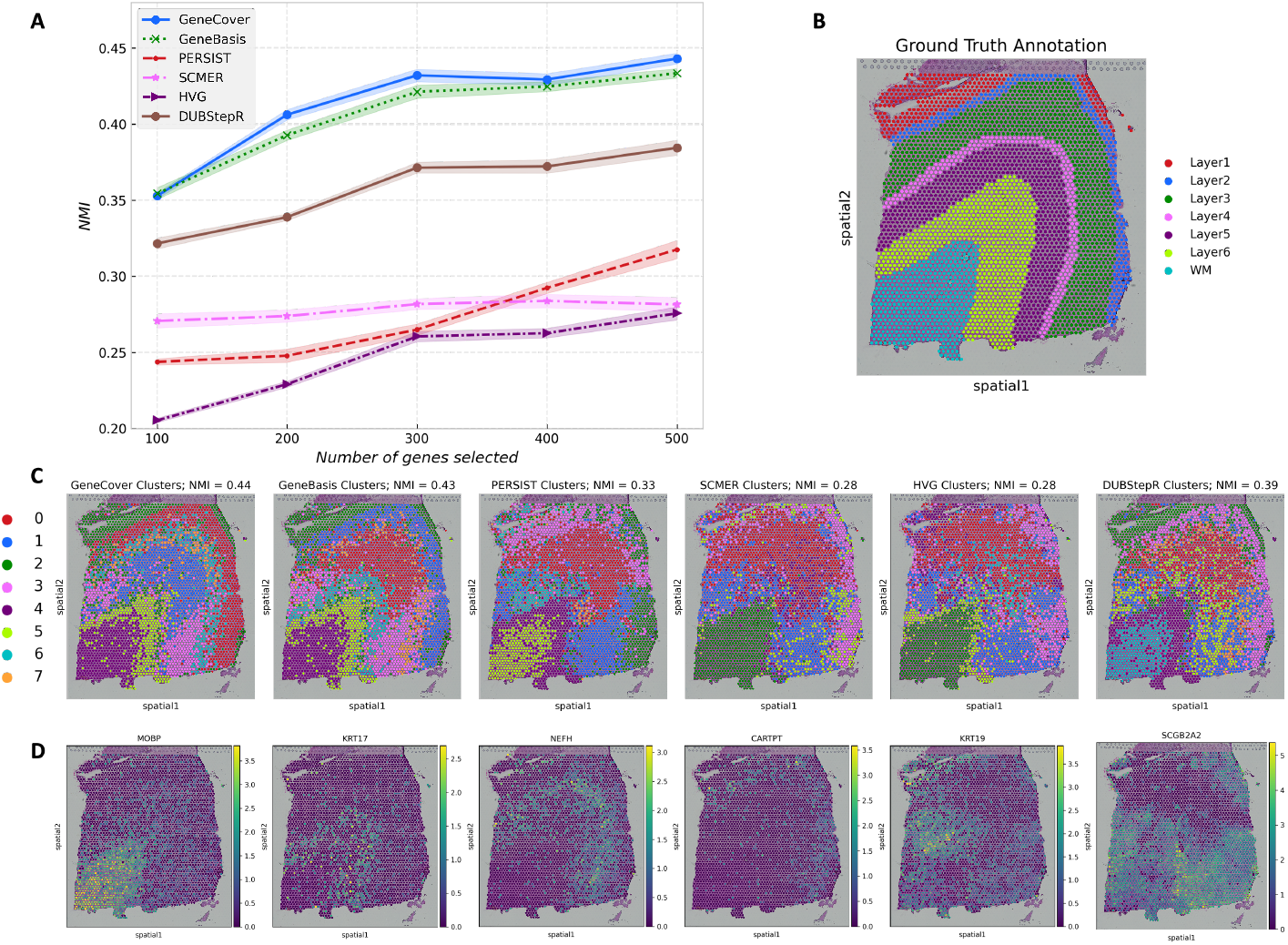
Benchmarking performance of geneCover and competing methods on the DLPFC dataset. To obtain the geneCover marker panel, we apply Iterative-GeneCover in Algorithm 1 with parameters: {*k*_*t*_} _1:*T*_ = {100, 100, 100, 100, 100}, *m* = 3. (A) NMI versus the size of the marker gene panel for all label-free methods. (B) Manually annotated histological layers of the DLPFC. (C) Leiden clusters obtained using 500 genes selected by each method with the same random seed. (D) Expression of selected marker genes from geneCover.

Despite being a label-free method, geneCover is able to identify layer-enriched signals that closely align with the manually annotated layers in the DLPFC dataset. For example, in Figure 1D, the white matter region is predominantly defined by the geneCover marker MOBP. KRT17 shows higher expression in layer 6, while the elevated expression of NEFH marks the boundary between layer 3 and 4. Likewise, layer 3 is enriched for CARTPT, and KRT19 is highly expressed in layer 1. These findings highlight geneCover’s ability to detect biologically meaningful signals even without pre-defined labels. Additionally, as label-free marker gene selection methods like geneCover identify genes from all sources of variability, some genes may capture biological processes that are not directly associated with anatomical structures. For instance, we observe that SCGB2A2 is the most differentially expressed gene in Cluster 3. Interestingly, its expression shows spatial variability, even though it does not correspond to any of the annotated layers.

In summary, the benchmark analysis validates the effectiveness of geneCover in recovering biologically meaningful structures in spatial transcriptomics data. Its performance underscores its potential for providing more accurate and biologically relevant insights into additional spatial transcriptomic and scRNA-seq data.

### 3.2 GeneCover Improves Resolution in Single Cell and Spatial Transcriptomics Discovery

#### CBMC

In this section, we highlight how geneCover discovers a minimally redundant set of highly specific marker genes that characterize the diverse cell types in the CBMC dataset. When restricting the marker panel to 50 genes for each method, geneCover identifies a minimally redundant set of genes that effectively captures the diverse cellular architecture within the cord blood dataset. Figure 2A and 2C (Figure S1A, S1B, S1C) show the Spearman’s correlation matrices for the first 50 genes identified by geneBasis and geneCover (SCMER, PERSIST, DUBStepR), respectively, with genes reordered using hierarchical clustering. Notably, geneCover identifies a more distinct set of correlated gene groups, as indicated by more and clearer diagonal blocks in Figure 2A compared to Figure 2C. Moreover, the correlated gene groups identified by geneCover are visibly smaller, indicating its ability to reduce redundancy through the minimal set-covering approach. We also observe that geneCover selects significantly fewer redundant marker genes for certain cell populations. For example, while SCMER, PERSIST, DUBStepR, and geneBasis identify multiple highly correlated markers for the mouse cell population, geneCover selects only the MYL3 gene to represent this group. This highlights how geneCover efficiently explores the complex correlation structures within the omics data and selects a compact, non-redundant set of marker genes.

**Fig. 2:**
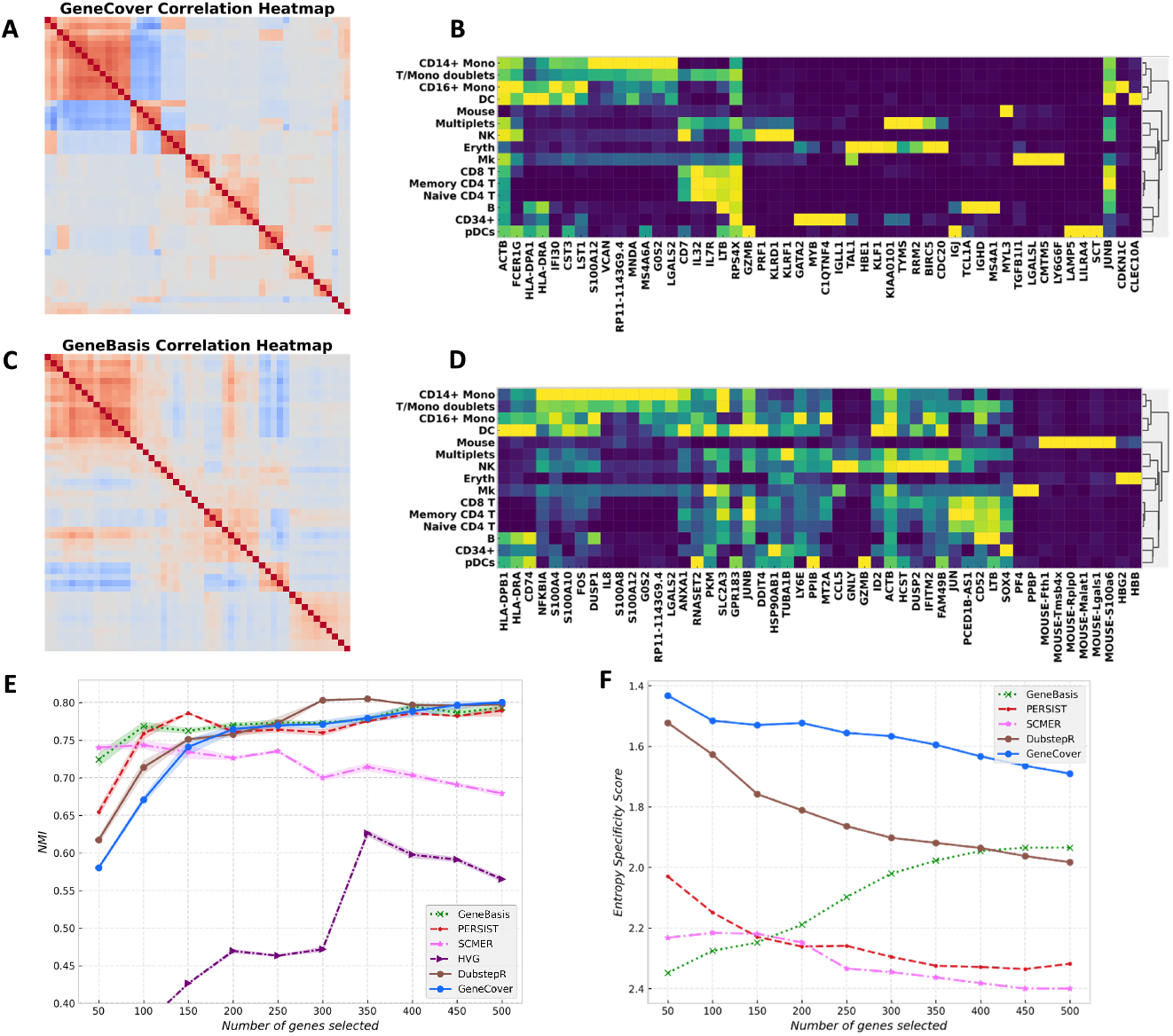
GeneCover identifies a minimally redundant set of marker genes characterizing diverse cell types in the CBMC dataset: To obtain the geneCover marker panel, we apply Iterative-GeneCover in Algorithm 1 with parameters: {*k*_*t*_}_1:*T*_ = {50, 50, 50, 50, 50, 50, 50, 50, 50, 50}, *m* = 6. (A) Spearman correlation heatmap of the 50 geneCover marker genes identified by geneCover, with gene reordered by hierarchical clustering. B) Expression matrix of geneCover markers, with same ordering as in (A), in cell types. Gene expression is standardized to [0, 1] range. The color intensity represents the level of the normalized expression. (C) Same as (A) but for geneBasis markers. (D) Same as (B) but for geneBasis markers. (E) NMI versus marker panel size for different label-free methods. (F) Specificity score versus marker panel size for different label-free methods. Lower score indicate better marker specificity.

With the ability to capture diverse cellular landscapes, geneCover markers can effectively resolve cell states within the CBMC dataset. Using a similar comparison procedure as in the previous subsection, we find that geneCover aligns with the top-performing methods in recovering cell types, as shown in Figure 2E. The normalized mutual information (NMI) scores for geneCover grow steadily with the marker panel sizes, and they are comparable to geneBasis, DUBStepR, and PERSIST.

Although these methods achieve similar performance in cell type recovery, they exhibit notable differences in marker gene characteristics. Figure 2F illustrates that geneCover markers are the most specific to individual cell types among all imputation-based methods across all marker panel sizes based on the specificity score (See Supplementary Section 2), while the markers from other imputation-based methods are more broadly expressed across multiple cell types but less informative about the cell types. To illustrate, in the size-50 panel, geneCover identifies CDKN1C as uniquely characterizing the CD16+ Mono cell type, and CLEC10A is exclusively expressed in DC cells. In contrast, the markers identified by geneBasis tend to favor widely expressed genes with broader variation patterns, as demonstrated by the diffused cell-type expression pattern in the matrix plots (Figure 2D). Although this selection may improve clustering accuracy—since geneBasis achieves strong performance with 50 markers (as shown in Figure 2E)—it does not summarize a rich portfolio of cell-type-specific expression patterns, potentially limiting deeper biological insights.

Notably, geneCover is also capable of distinguishing hierarchical gene expression patterns. For example, as demonstrated in Figure 2B, KIAA0101 is expressed in both the multiples and erythroid cell populations, while KLF1 is uniquely expressed in erythroid cells. Even though KIAA0101 encompasses the expression pattern of KLF1, geneCover is still able to distinguish these two gene groups with overlapping yet distinct correlation structures.

#### Mouse Brain Visium HD

We demonstrate that geneCover can effectively resolve hippocampal subfields in the mouse brain. We selected 200 marker genes using geneCover, geneBasis, and DUBStepR and applied the Leiden clustering algorithm on a cell-neighborhood graph, which was generated from the expression profile restricted to these marker genes (Figure 3A, 3B). Here, we avoid using principal components, which could downplay the contribution of marker genes that characterize spatial organization with very low abundance. As a comparison, we also applied the conventional clustering pipeline using 200 principal components of the entire transcriptome, followed by Leiden clustering (Figure 3C). To ensure a fair comparison, we adjusted the Leiden clustering resolution for each method so that all pipelines produced 15 clusters, matching the number of clusters provided by 10X Genomics [15]. The resolution parameter controls the coarseness of clustering, where higher values lead to finer subdivisions and more clusters. The average resolutions over five random seeds were 0.99 for geneCover, 1.62 for geneBasis, 1.30 for DUBStepR, and 1.06 for the conventional pipeline. Notably, geneCover required the lowest resolution to achieve 15 clusters, which may suggest that the selected gene panel captures a diverse range of expression patterns, facilitating natural partitioning of the data.

**Fig. 3:**
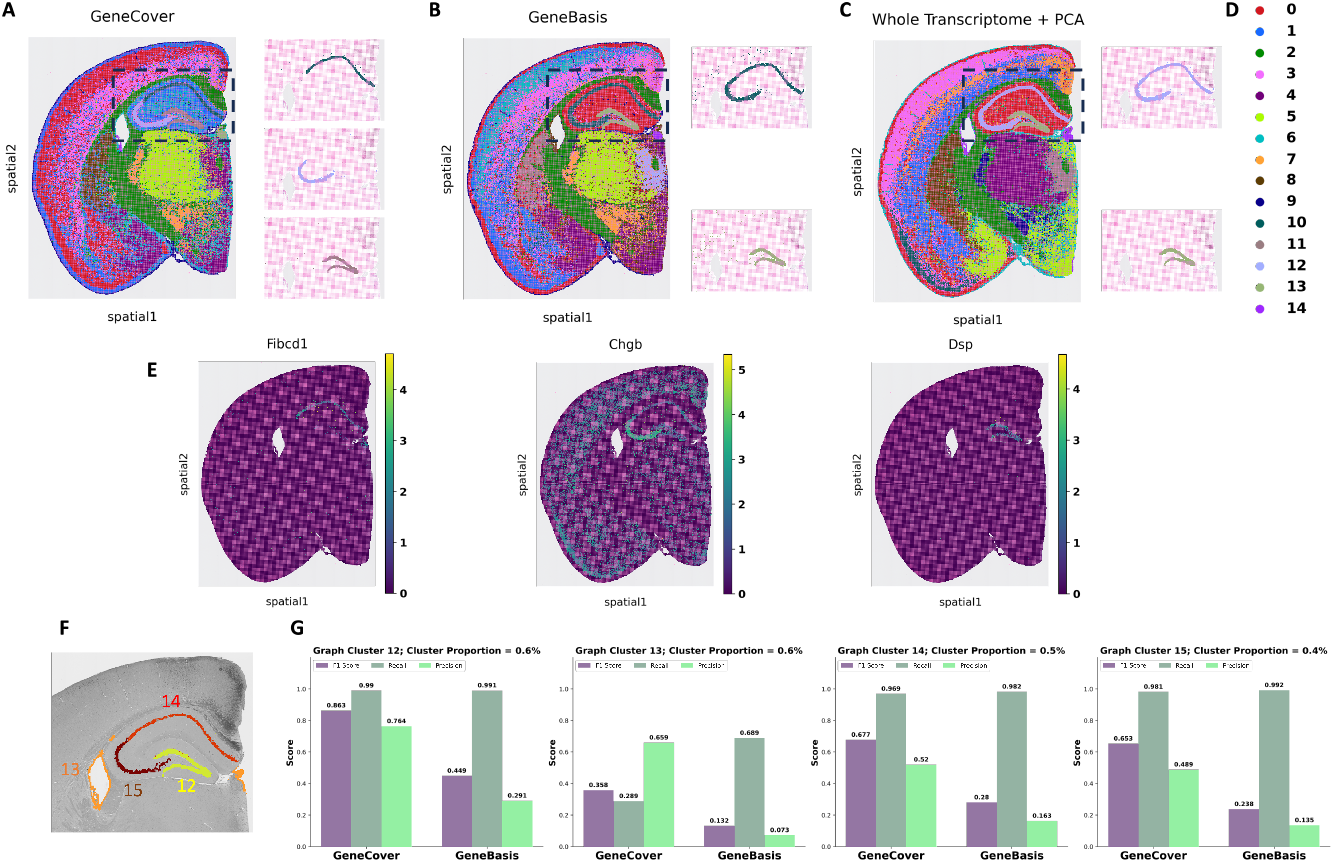
GeneCover resolves hippocampal subfields in the mouse brain: To obtain the geneCover marker panel, we apply Iterative-GeneCover in Algorithm 1 with parameters: {*k*_*t*_}_1:*T*_ = {80, 60, 60}, *m* = 3. (A) Leiden clusters learned from 200 geneCover markers, with a highlight of clusters in the CA1-CA3 subiculum and dentate gyrus regions. (B) Same as Panel A but for 200 geneBasis markers. (C) Same as Panel A but for 200 principal components from the whole transcriptome. (D) Legend for clusters in Panel A, B, and C. (E) Differentially expressed geneCover markers for geneCover clusters 10, 11, and 12. (F) Four graph-based clusters provided by 10x Genomics with the lowest cell abundance. (G) Comparison of geneCover and geneBasis in resolving rare spatial organization in the mouse brain.

While all methods manage to identify the dentate gyrus in the mouse brain, geneCover uniquely divides the CA1-CA3 subiculum into two distinct regions (Figure 3A & Figure S2A), while none of geneBasis, whole-transcriptome + PCA, and DUBStepR is capable of resolving this important hippocampal subregion, regardless of the random seeds used for clustering (Figure S2B, S2C, S2D). Even with increased clustering resolution, geneBasis and DUBStepR markers still struggle to segregate the area (Figure S3). This distinction is significant as the CA1-CA3 subiculum plays a crucial role in hippocampal function, contributing to memory formation and spatial navigation. Importantly, the two clusters identified by geneCover are transcriptionally distinct, as evidenced by the marker genes within the geneCover panel (Figure 3E). For example, FIBCD1 expression is uniquely localized to geneCover cluster 10 (Figure 3A, top right), which corresponds to the first division of the CA1-CA3 subiculum, while CHGB shows the highest expression in cluster 12 (Figure 3A, middle right), representing the second division of the CA1-CA3 subiculum. Region-specific signals representing the division of CA1-CA3 are primarily detected by geneCover, while geneBasis and DUBStepR fail to capture many of these signals. We selected multiple marker genes differentially expressed between the two divisions: FIBCD1, SPINK8, IQGAP2, and LEFTY1 for the first division, and CHGB, DNM1, SLC7A7, and SNAP25 for the second division. Based on Figure S4 (Row 1-3), geneBasis fails to detect any markers for the first division, while DUBStepR identifies only FIBCD1. In contrast, geneCover successfully captures FIBCD1, SPINK8 and IQGAP2. For the second division, although all three methods identify DNM1, SLC7A7, and SNAP25, the most region-specific signal, CHGB, is uniquely detected by geneCover. The abundance of these region-specific markers in the geneCover panel facilitates the identification of these highly refined spatial organizations.

To further quantify how well geneCover resolves these delicate spatial organizations, we compared its performance to geneBasis using the 15 graph-based clusters provided by 10X Genomics (Figure S5) as a reference. We focused specifically on four reference clusters with the smallest cell abundances (clusters 12–15 in Figure 3F), each comprising less than 1% of the total cell population. We matched clusters from geneCover and geneBasis to the 10X Genomics clusters using the F1 score (See Supplementary Section 3 for details). According to Figure 3G, the clusters identified by geneCover consistently demonstrate better matching qualities with the four rarest reference 10X Genomics clusters based on F1 score (See Figure S6 for matching quality comparisons on all 10X Genomics clusters). This result highlights geneCover’s ability to enhance the resolution of spatial transcriptomics discovery, particularly in identifying highly refined spatial organizations, using a compact and minimally redundant set of genes.

#### scFFPE Breast Cancer

GeneCover markers facilitate the identification of a transcriptionally distinct immune cell subpopulation that was absent from the original cell type annotations in the scFFPE breast cancer dataset. Specifically, within the originally labeled Macrophage 1 cell population (Figure 4A), geneCover uniquely identifies a subpopulation (Cluster 12 in Figure 4C) using 300 markers, a distinction that geneBasis struggles to achieve (Figure S7). Moreover, we demonstrate that geneCover can reliably identify this potential immune subpopulation even with marker panels reduced to 100 or 200 genes. This performance remains robust across different random seeds used for clustering (Figure S8).

**Fig. 4:**
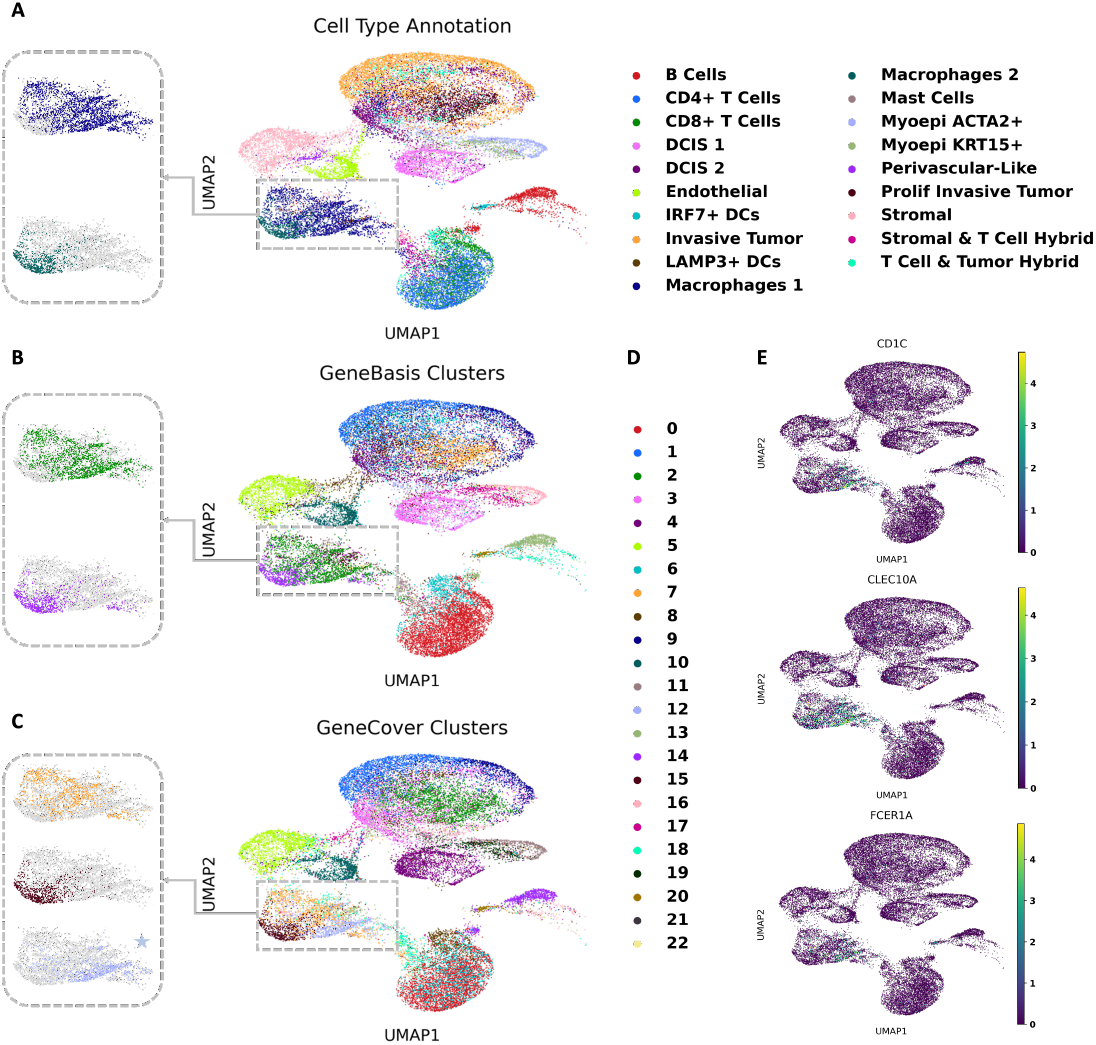
GeneCover Markers Facilitate the Identification of a Transcriptionally Distinct Immune Cell Subpopulation in Breast Cancer: To obtain the geneCover marker panel, we apply Iterative-GeneCover in Algorithm 1 with parameters: {*k*_*t*_}_1:*T*_ = {100, 100, 100}, *m* = 3. (A) UMAP visualization of cell type annotations provided by the dataset, with a zoom-in on the Macrophage 1 and Macrophage 2 subpopulations. (B) Data-driven Leiden clusters learned from 300 geneBasis markers using the standard pipeline. (C) Same as Panel B, but for 300 geneCover markers. (D) Legend for the clusters in Panels B and C. (E) Differentially expressed genes for geneCover Cluster 12.

Based on differential expression analysis of geneCover cluster 12, we hypothesize that this immune cell population may be related to dendritic cells, as geneCover identifies CD1C, CLEC10A, and FCER1A, which are all well-established markers of dendritic cells, as marker genes for this cluster (Figure 4E). CD1C is a marker of conventional dendritic cells type 2 (cDC2), which are crucial for presenting antigens and initiating immune responses. CLEC10A is specifically expressed on CD1C+ dendritic cells, enhancing their cytokine secretion in response to toll-like receptor stimulation, which contributes to their role in immune surveillance [23]. Notably, CLEC10A is also identified by geneCover in the CBMC dataset, where it is exclusively expressed in dendritic cells (Figure 2B, the last gene). Lastly, FCER1A encodes the alpha chain of the high-affinity IgE receptor, which is expressed on dendritic cells and plays a critical role in mediating allergic responses by promoting antigen presentation and activation of immune cells in response to IgE-bound allergens [24]. Together, the identification of these marker genes suggests that the transcriptionally distinct immune cell subpopulation uncovered by geneCover may represent a previously unrecognized subset of dendritic cells within the tumor microenvironment.

The ability of geneCover to enhance the resolution of omics data analysis offers the potential for novel hypotheses regarding subtle cell types, shedding light on cell populations that may have been overlooked in previous studies.

### 3.3 Scalability

In this section, we demonstrate the scalability of geneCover when applied to large omics datasets. As shown in Table 1, geneCover significantly outperforms other label-free marker gene selection methods in terms of run time across the four datasets of consideration. Here, we include only label-free methods that allow the specification of marker panel size.

**Table 1:**
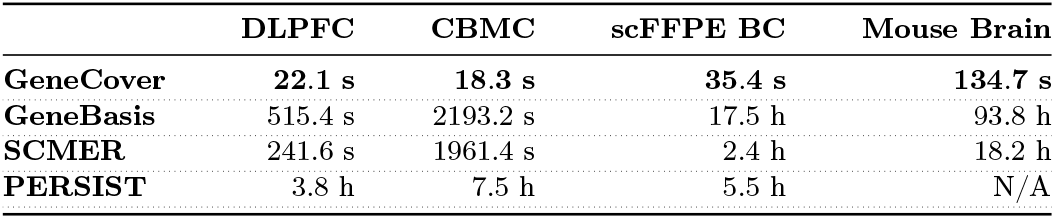
Empirical Run Time of label-free Marker Gene Selection Methods on Omics Datasets: The time is measured in either seconds (s) or hours (h). For each method, we obtained 100 markers for the DLPFC and CBMC datasets, 300 markers for the scFFPE breast cancer dataset ({*k*_*t*_}_1:*T*_ = [100, 100, 100], *m* = 3 for geneCover), and 200 markers for the mouse brain Visium HD dataset ({*k*_*t*_}_1:*T*_ = [80, 60, 60], *m* = 3 for geneCover). GeneCover and geneBasis were run using the Intel Core i9 13900K CPU, while SCMER and PERSIST were run on the NVIDIA GeForce RTX 4090 GPU. N/A indicates memory overflow.

|  | DLPFC | CBMC | scFFPE BC | Mouse Brain |
| --- | --- | --- | --- | --- |
| **GeneCover** | **22.1 s** | **18.3 s** | **35.4 s** | **134.7 s** |
| **GeneBasis** | 515.4 s | 2193.2 s | 17.5 h | 93.8 h |
| **SCMER** | 241.6 s | 1961.4 s | 2.4 h | 18.2 h |
| **PERSIST** | 3.8 h | 7.5 h | 5.5 h | N/A |

In particular, for the mouse brain Visium HD dataset, which contains approximately 100,000 bins, geneCover completes its task in just 134.7 seconds, making it approximately 500 times faster than the runner-up, SCMER, which requires 18.2 hours. Additionally, geneBasis takes over 93 hours to generate the marker panel, and PERSIST is unable to handle the dataset due to memory overflow, regardless of the various batch sizes tested. Furthermore, iterative geneCover provides a scalable solution even when a much larger gene panel size is required. To illustrate, we applied the algorithm to all four datasets and gradually increased the gene panel size by 100 at a time, reaching a final panel size of 1000 genes. As shown in Figure S9, geneCover returns the solution in less than 3 minutes for all four datasets.

The observed scalability of geneCover can be attributed to its focus on gene-gene correlations. The most computationally intensive step is the calculation of the correlation matrix, which scales with the number of cells. However, this step can typically be executed efficiently using parallel computing. More importantly, because the input dimension for the minimal set covering problem is determined solely by the number of genes, the run time of the set covering algorithm remains invariant to the number of cells and depends only on the number of genes being considered. In contrast, the three imputation-based methods have time complexities that scale with both the number of cells and the number of genes. As a result, their run times increase considerably as the dataset size grows, making them significantly slower on large-scale datasets.

However, we acknowledge that geneCover’s runtime efficiency may be compromised when the required (incremental) marker panel size, *k*_*t*_, is too large. This limitation arises fundamentally from the reduction of the correlation threshold λ as *k*_*t*_ increases (Figure S10). For instance, in the DLPFC dataset, when λ falls below 0.09 — which corresponds to requiring *k*_*t*_ > 1600 in a single run of geneCover — the algorithm is no longer able to return the optimal solution within a few seconds. This occurs because a lower λ leads to an increase in gene neighborhood size, thereby expanding the coverage capacity of all genes. Consequently, the number of feasible covering sets may grow significantly, prolonging the search for an optimal solution. Furthermore, a lower λ introduces greater overlap among gene neighborhoods, which can obscure the distinction of the genome-wide correlation structure. To address this, if a large gene panel is required, we recommend using iterative geneCover with a properly chosen step size to gradually expand the gene set. Compared to obtaining all selected genes in a single run, this approach not only ensures efficient runtime but also enables genes to be sequentially selected from distinct correlated gene groups, thereby capturing diverse signals that characterize the cellular landscape.

As advances in high-throughput whole-transcriptome spatial transcriptomics continue to push cellular resolution to new levels, the scalability of marker gene selection methods becomes increasingly critical. geneCover achieves excellent runtime efficiency across large-scale datasets. While selecting extremely large gene panels in a single run may increase computational demands due to a lowered correlation threshold λ, this can be effectively addressed using the proposed iterative approach. By leveraging its ability to sequentially expand gene sets, iterative geneCover ensures robust and efficient identification of marker gene panels of varying sizes, making it well-suited for evolving spatial transcriptomics technologies.

### 3.4 Robustness to Hyperparameter Selection

To assess the robustness of geneCover’s ability to resolve fine-grained spatial organizations, we examined the impact of hyperparameter selection on its clustering performance of the Mouse brain Visium HD dataset [15]. Specifically, we evaluated how variations in key parameters—the gene neighborhood size threshold (*m*) and the sequence of incremental sizes ({*k*_*t* 1:*T*_})—affect the resolution of spatial structures. First, we fixed {*k*_*t*_}_1:*T*_ = {80, 60, 60} (as used in the previous analysis) and gradually increased *m*. According to Figure S4 (Rows 4–9), despite requiring each selected marker to cover more genes, the geneCover marker panel still retains a substantial number of region-specific signals that characterize the two CA1-CA3 subdivisions, allowing these highly refined regions to remain identifiable by Leiden clustering even with *m* = 30 (Figure S11A). This seemingly counterintuitive result can be explained by the self-adjusting nature of the algorithm: given the fixed {*k*_*t*_}_1:*T*_, as *m* increases, the final correlation threshold λ determined by the binary search decreases (Figure S12), allowing each selected marker to cover a broader set of genes. Thus, even though a stricter coverage requirement is imposed, the increase in gene neighborhood sizes ensures that genes capturing fine-grained spatial structures will be preserved in the covering panel.

Secondly, we fixed *m* = 3 and varied {*k*_*t*_}_1:*T*_ to generate a geneCover panel of size 200. We found that the CA1–CA3 subregion could be successfully segregated using a geneCover panel generated with fewer iterations (Figure S11B, {*k*_*t*_}_1:*T*_ = [100, 100]). However, as *T* increases beyond 3, Leiden clustering fails to detect this division (Figure S11B, {*k*_*t*_}_1:*T*_ = [50, 50, 50, 50], {*k*_*t*_}_1:*T*_ = [40, 40, 40, 40, 40]). This behavior arises because larger *T* values require smaller *k*_*t*_, increasing the final λ at each iteration. As a result, the selected markers exhibit strong correlations with the genes they cover, and they primarily capture broad variability patterns rather than finer-scale biological distinctions. This suggests that small-size coverings may not include enough highly specific genes crucial for detecting certain subregion differences. Indeed, as shown in Figure S4 (Row 11-12), increasing *T* while fixing the total gene panel size results in the loss of some differentially expressed genes that define the CA1–CA3 segregation, particularly CHGB, which characterizes the second division of CA1–CA3. However, we observe that these lost signals can be restored by expanding the marker panel beyond the fixed size of 200 through additional iterations. For example, with one additional iteration ({*k*_*t*_}_1:*T*_ = [50, 50, 50, 50, 50]), selected differentially expressed genes distinguishing the two subdivisions—FIBCD1, SPINK8, IQGAP2, LEFTY1 for the first division and CHGB, DNM1, SLC17A7, SNAP25 for the second—are all successfully identified by geneCover (Figure S4, Row 13). Similarly, setting *k*_*t*_ = 40 for *t* = 1, …, 6 to extend the panel to 240 genes restores FIBCD1 and SPINK8, as well as CHGB in the expanded panel (Figure S4, Row 14).

To summarize, across a range of reasonable hyperparameter choices, geneCover retains a substantial proportion of region-specific signals, enabling highly refined tissue organization to remain identifiable by Leiden clustering. However, we find that overly fragmented coverings (small *k*_*t*_) may limit the resolution of biological signals under our clustering pipeline. This effect is naturally mitigated as the panel expands with additional iterations, allowing geneCover to sequentially incorporate a broader set of correlation structures within the transcriptome.

### 3.5 Marker Gene Selection Across Samples

To identify robust biological signals that characterize conserved cellular programs, we can naturally extend geneCover to enable marker gene selection across multiple samples. In this generalized formulation, we require every gene in the transcriptome *G* to be covered by at least one of the selected marker genes in all samples.

Let *B* denote the set of samples, and let 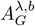 be the binary adjacency matrix determined by gene-gene correlations within each sample *b* ∈ *B*. The generalized integer programming formulation is given by:

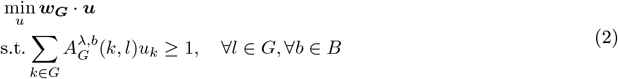

where the objective minimizes the weighted selection of marker genes while ensuring that each gene *l* in the transcriptome is covered in every sample (See detail in Algorithm S1). Notably, the original integer programming formulation defined in formulation 1 is a special case of this generalized formulation, where *B* contains a single sample.

We applied the generalized geneCover to extract marker genes that characterize shared tissue organization of the human dorsolateral prefrontal cortex across three DLPFC samples — #151507, #151673, and #1516569 — collected from different donors (Figure 5A). As shown in Figure 5B, a subset of the selected marker genes — MOBP, KRT17, PCP4, NEFH, CARTPT, HPCAL1, and VIM — exhibits strong layer-specific expression patterns, covering Layers 1 through 6 and the white matter. Figure 5C–E further confirms that the expression of each gene is concentrated within distinct laminar regions across all three tissue sections, providing evidence that these genes robustly mark their respective layers.

**Fig. 5:**
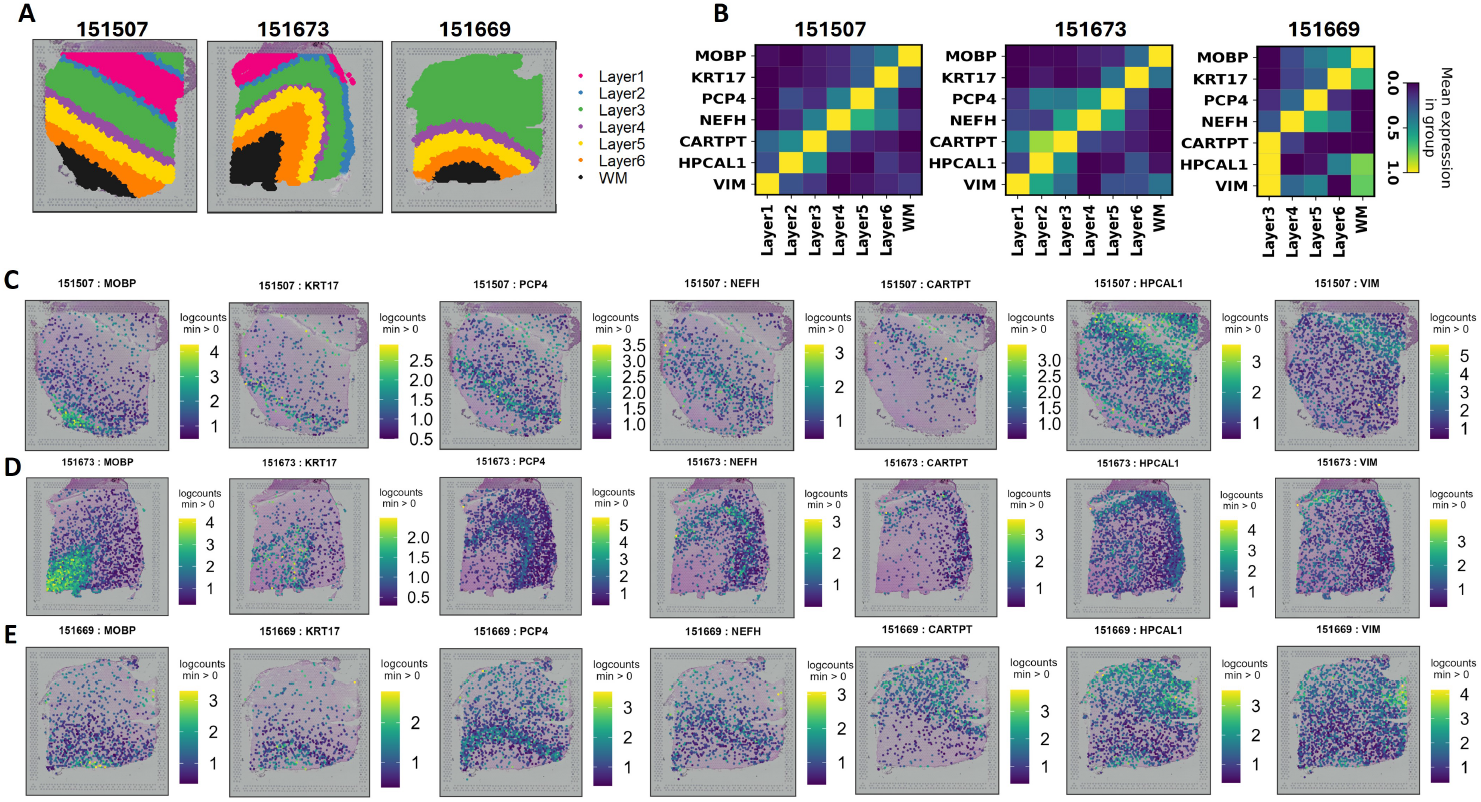
GeneCover Marker Selection Across Three Samples of the DLPFC Dataset: We apply Algorithm S1 with {*k*_*t*_}_1:*T*_ = {100, 100, 100}, *m* = 3. (A) Annotated histological layers of three DLPFC tissue samples. (B) Normalized expression matrix of the selected geneCover markers across different layers in each sample. (C) Spatial expression (in log counts) patterns of the selected markers in sample #151507. (D) Same as (C) for sample #151673. (E) Same as (C) for sample #151669.

Additionally, their spatial expression patterns illustrate a progressive transition across cortical layers. According to 5C–E, from left to right, the regions with predominant expression shift from the white matter to the outer cortical layers, reflecting a structured gradient in the human cortex. Notably, although Layers 1 and 2 are absent in sample #151669, CARTPT, HPCAL1, and VIM remain spatially localized within different regions of Layer 3, suggesting potential intra-layer heterogeneity in this donor sample.

Overall, geneCover identifies coherent layer-specific gene expression patterns that are conserved across all three donors, underscoring its ability to detect shared transcriptional signatures.

## 4 Discussion

In this work, we propose geneCover as a novel label-free marker gene selection algorithm for single-cell and spatial transcriptomics. GeneCover employs a minimal-weight set covering approach to identify a minimally redundant panel of marker genes that represent distinct correlation structures within the transcriptome. Our study demonstrates that geneCover provides a robust and scalable solution for label-free marker gene selection, outperforming existing label-free methods in both computational efficiency and accuracy in resolving refined spatial organization or rare cell types in spatial transcriptomics and scRNA-seq data.

GeneCover excels at identifying a minimally redundant marker panel that captures various sources of meaningful transcriptional variability. By leveraging minimal set covering to explore distinct gene-gene correlation groups, geneCover facilitates the discovery of granular biological signals as effectively as it identifies marker genes with large variations across cell types. This allows geneCover to significantly enhance the resolution of scRNA-seq and spatial transcriptomics discovery. Notably, geneCover markers enable the division of the highly refined CA1-CA3 subiculum in the mouse brain hippocampus into two distinct regions—a distinction that other existing strategies fail to achieve. In addition, using geneCover markers, we identify a transcriptionally distinct immune cell subpopulation characterized by dendritic cell markers, suggesting the potential to uncover previously unrecognized cell types or subpopulations. The ability to identify finely organized and transcriptionally distinct cell subpopulations is crucial for advancing our understanding of tissue heterogeneity, disease progression, and cellular dynamics.

However, we note that the conventional clustering pipeline we use introduces certain limitations that may impact the identification of rare cell states. In all of our analyses, we apply Leiden clustering either on the PCA-transformed marker gene expression data or directly on the marker gene expression matrix. This approach has two main constraints. First, principal components primarily capture dominant cell-type signals from populations characterized by many highly expressed, variable marker genes, which may result in the loss of signals defining rare cell states. Second, even when PCA is omitted, graph-based clustering methods typically construct the cell-cell graph using Euclidean distance on expression vectors. This metric can be insensitive to highly specific marker genes that distinguish subtle cell populations, potentially affecting their detection. Despite these challenges, geneCover markers still enhance resolution in identifying rare cell populations. This reflects the ability of our iterative covering framework to identify an adequate number of rare cell-type signals that remain recognizable by conventional clustering pipelines. Future work could address these limitations by incorporating dimensionality reduction techniques that better capture local signals characterizing subtle cell populations [25], alongside specialized rare cell discovery methods [26, 27, 28, 29].

Beyond its improved resolution, geneCover achieves significantly faster empirical run times compared to other label-free marker gene selection methods, particularly on large omics datasets. As cellular resolution continues to expand in whole-transcriptome spatial transcriptomics, the ability to process larger and more complex datasets efficiently will be critical. GeneCover offers a highly practical solution for modern highthroughput analyses and is well-positioned to adapt to these growing data modalities.

Furthermore, our analysis demonstrates that geneCover is fairly robust to hyperparameter selection. We show that adjusting the gene neighborhood size threshold *m* and the sequence of incremental sizes {*k*_*t*_}_1:*T*_ allows for fine-tuning marker gene selection while preserving signals necessary to identify spatially refined structures. One of the key findings is the self-adjusting property of geneCover: for a fixed marker panel size, increasing *m* lowers the final correlation threshold λ, ensuring that marker genes remain effective in capturing fine-grained spatial structures even under stricter coverage constraints. However, lowering *k*_*t*_ excessively without compensatory adjustments can lead to a loss of resolution due to an increase in λ, which results in a bias toward selecting genes that capture broad variability rather than region-specific signals. To capture additional signals defining rare cell identities, the number of iterations *T* can be increased, allowing the panel to expand beyond the previously fixed size.

Based on these observations, we provide the following general recommendations for selecting geneCover’s hyperparameters. For *m*, we suggest starting with a moderate value (e.g., *m* ≤ 6) to prevent over-filtering of signals even though geneCover is relatively robust to this parameter. For the incremental sizes {*k*_*t*_}_1:*T*_, we recommend using a balanced sequence with moderate values of *k*_*t*_, such as {100, 100, …}, to maintain consistency in marker selection while ensuring comprehensive coverage of λ. The general principle is to finetune hyperparameters so that the final correlation threshold λ effectively distinguishes biologically relevant correlated gene groups. Since geneCover provides functionality to monitor λ in the package, users can empirically determine suitable values based on their dataset. Our proposed iterative approach also naturally supports this tuning process, as each iteration yields a different λ. This allows us to sequentially explore signals representing the diverse correlation structure of the transcriptome.

Lastly, we demonstrate that geneCover can be naturally extended to identify shared molecular signals across multiple tissue samples, ensuring consistent selection of marker genes that may facilitate the discovery of conserved biological structures. Applying this framework to three DLPFC donor samples, we show that jointly identified marker genes accurately reflect the cortical layer organization with highly localized expression patterns. These markers not only delineate Layers 1 through 6 and the white matter, but also exhibit structured spatial transitions across cortical layers. Additionally, subtle intra-layer heterogeneity observed in Layer 3 of donor sample #151669 highlights geneCover’s ability to detect variation within conserved spatial structures. With the expanding capacity of spatial transcriptomics technologies to encompass more tissue samples, our method provides a practical and scalable solution to investigate conserved transcriptional programs through shared molecular signatures that are critical for understanding key biological processes.

## Code Availability

The Python implementation of geneCover is publicly available on GitHub at: https://github.com/ANWANGJHU/GeneCover

## Acknowledgments

The work is supported by NSF Award 2124230. We would like to express our gratitude to Jean Fan, Luigi Marchionni, and Caleb Hallinan for their valuable discussions on dataset selection and computational experiment design. We are also grateful to Hongyu Cheng for insightful discussions on integer programming.

## Footnotes

1 ⟦*d*⟧ represents the whole transcriptome, and *G* is the index set of remaining genes after filtering the gene set.

