## Supplementary Material for "GeneCover: A Combinatorial Approach for Label-free Marker Gene Selection"


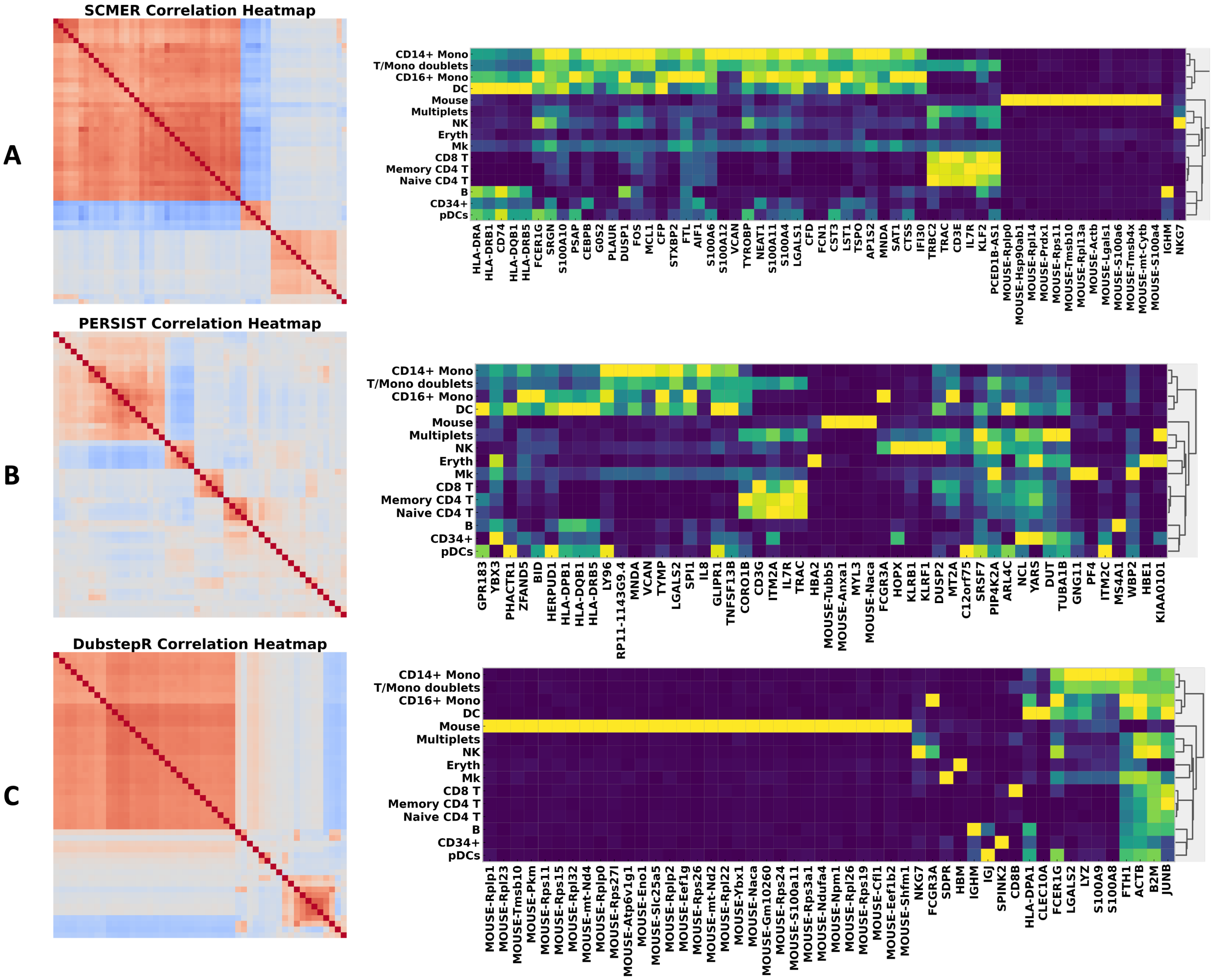


**Fig. 1.** (A) Left: Spearman correlation heatmap of the 50 SCMER marker genes, with gene reordered by hierarchical clustering; Right: Expression matrix of SCMER markers, with same ordering, in cell types. Gene expression is standardized to $[0, 1]$ range. The color intensity represents the level of the normalized expression. (B) Same as Panel A but for PERSIST markers. (C) Same as Panel A but for DUBStepR markers.

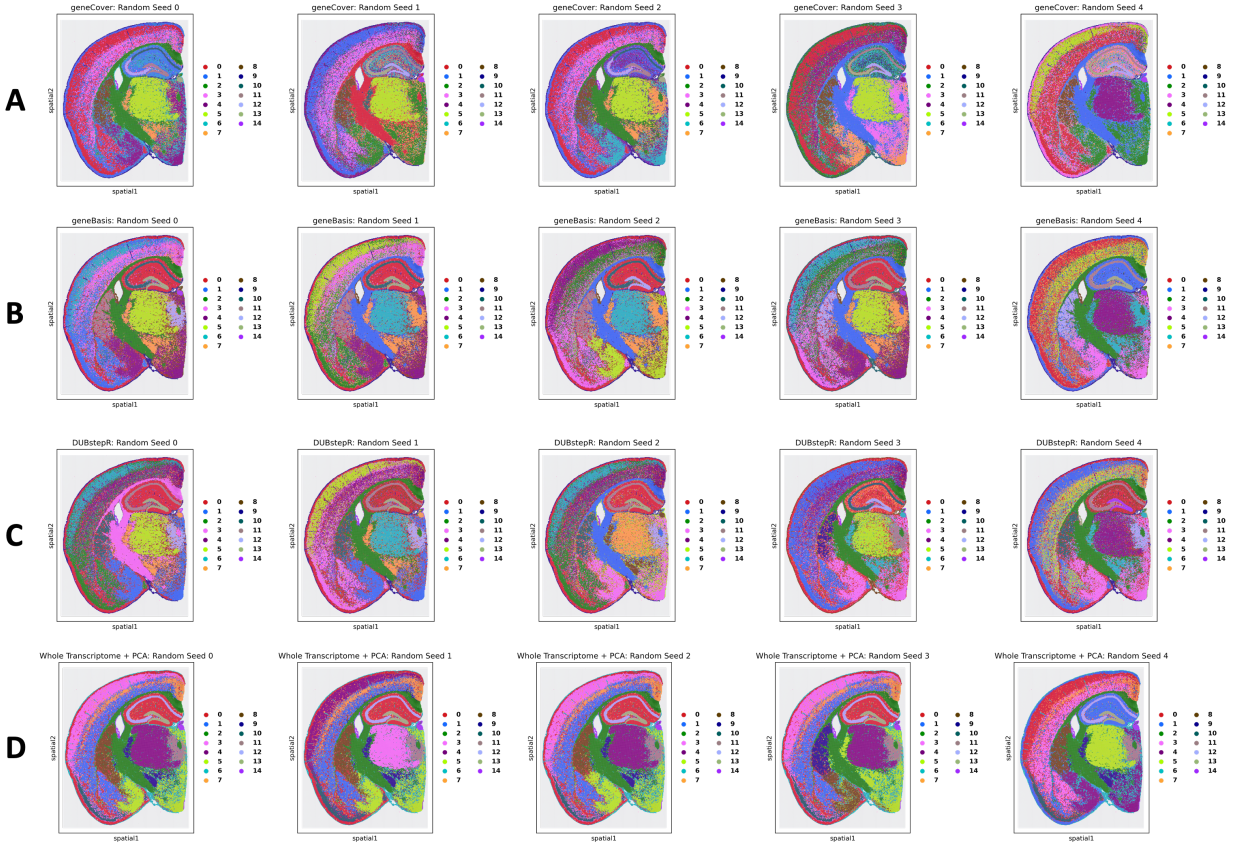


**Fig. 2. Leiden Clusters Over Random Seeds:** Leiden clusters obtained using 200 markers from geneCover, geneBasis, and DubstepR where we use the gene expressions to construct the cell neighborhood graph. The same clustering procedure is applied to 200 principal components of the whole transcriptome. (A) Leiden clusters learned from geneCover markers over five random seeds of the clustering algorithm. Clustering resolutions over random seeds: $[0.95, 1, 1, 1, 1.02]$. (B) Same as A but for geneBasis. Clustering resolutions over random seeds: $[1.6, 1.7, 1.595, 1.62, 1.6]$. (C) Same as A but for DUBStepR. Clustering resolutions over random seeds: $[1.3, 1.2, 1.32, 1.4, 1.3]$. (D) Same as A but for whole transcriptome data processed by PCA. Clustering resolutions over random seeds: $[1.1, 1.06, 1.05, 1.05, 1.05]$.

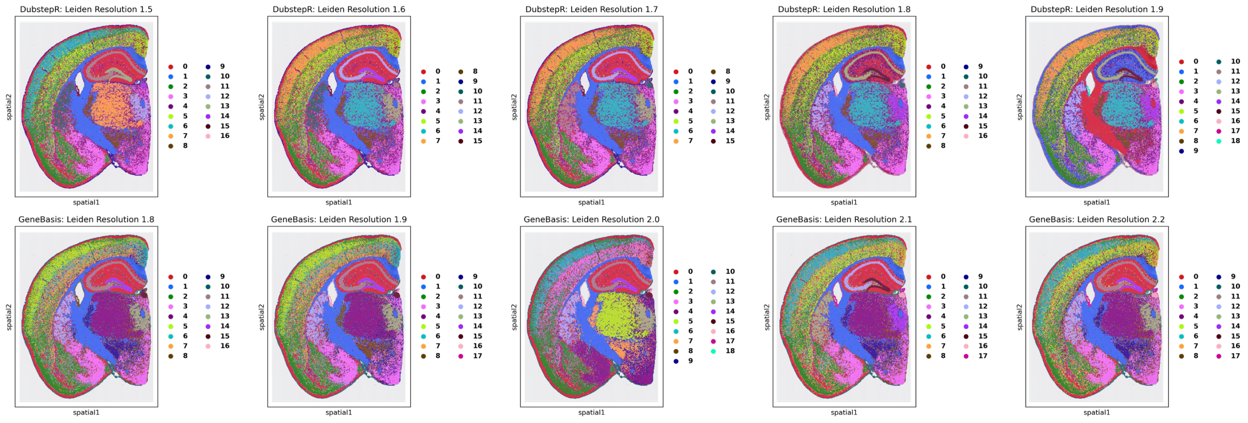


**Fig. 3. GeneBasis and DubstepR Leiden Clusters with Increased Clustering Resolution**

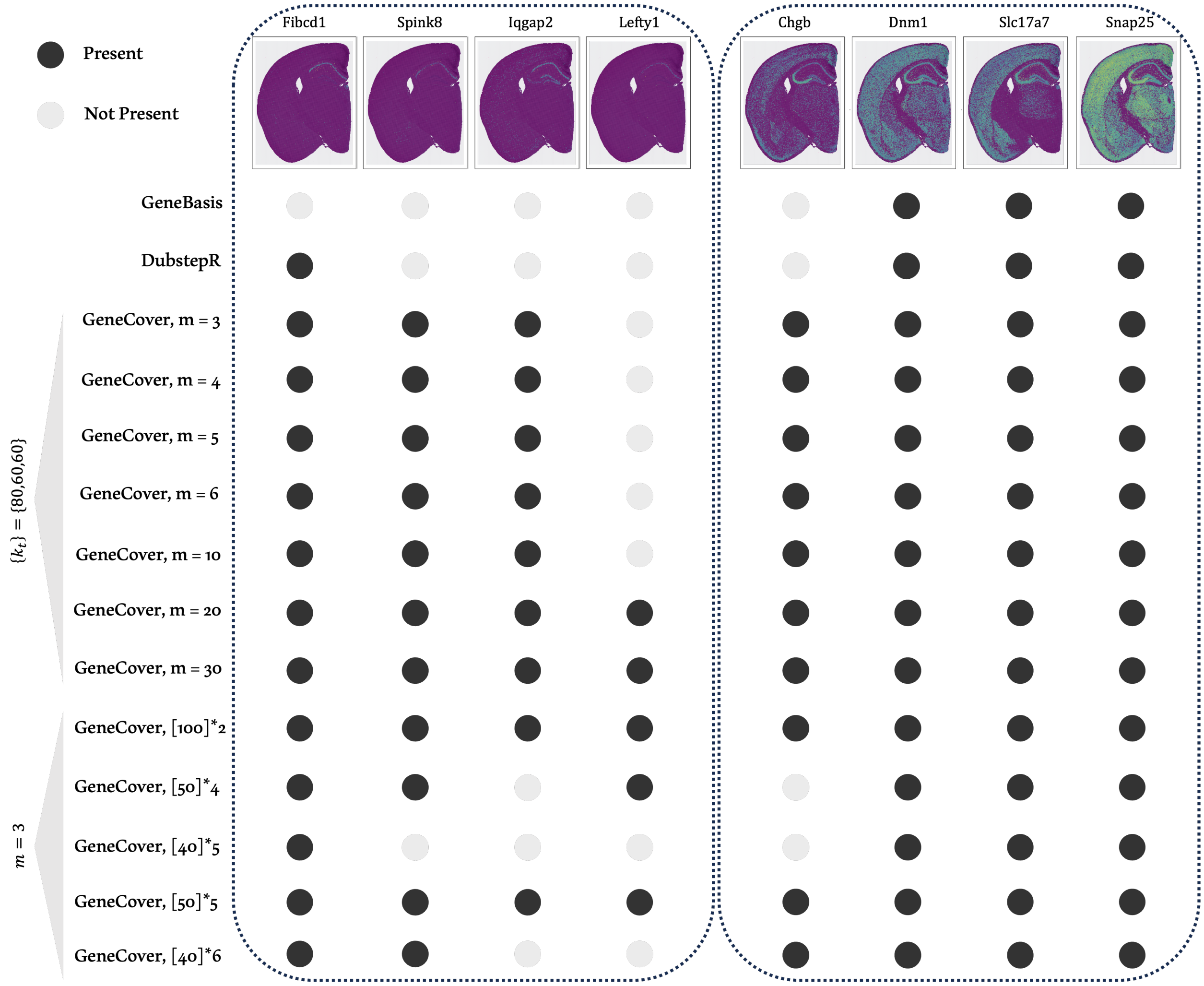


**Fig. 4. Top Differentially Expressed Genes in the Two Subdivisions of the CA1-CA3 Subiculum and Their Presence in Different Marker Panels.**

### 1 Dataset Processing and Marker Panel Generation

#### 1.1 Dataset Processing

**DLPFC** We used sample #151673 in DLPFC data. This dataset contains 3639 spots and 33538 gene expression measurements. We filter the genes that are expressed in less than 100 spots, normalize the counts per cell to 10000, log-transform the data, and then keep 10000 highly variable genes based on the dispersion score. We let $G$ to be the resulting gene sets.

**CBMC** The CBMC dataset contains 8617 cells with 20501 gene expression measurements and 10 protein measurements, where we only use the gene expressions for analysis. We filter genes that express in less than 30 cells but keep all the protein measurements. We then separately normalize gene counts per cell to 10000. Next, we log-transform the data. Eventually, we keep $\sim$ 8000 highly variable genes based on the dispersion score. We let $G$ to be the resulting gene sets. On this dataset, we find that running Leiden clustering directly on log-normalized marker gene expression data improves performance compared to using PCA-processed data for all label-free methods. As a result, we have opted not to use PCA-processed data in the NMI benchmark experiment for this dataset.

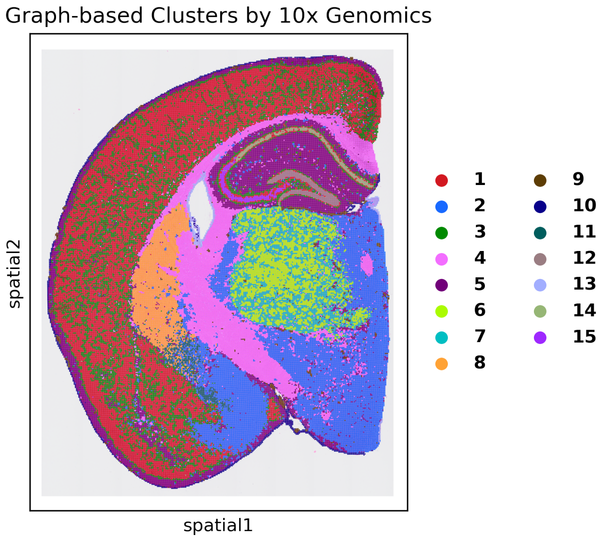


**Fig. 5. Graph-based Clusters From 10x Genomics** [1]

**Mouse Brain Visium HD** The Mouse Brain FFPE Visium HD dataset with 16 micron bin resolution contains 98917 bins and 19059 gene expression measurements. We filter out genes that are expressed in less than 100 bins and exclude cells that have no genes expressed in them. Next, we normalize the counts per cell to 10000 and log-transform the data. We observe that the highly variable gene feature selection will result in the removal of genes that characterize spatial organization with extremely low cell abundance, so we do not further subset the gene set. We let $G$ to be the resulting gene sets.

**scFFPE Breast Cancer** The scFFPE breast cancer scRNA-seq dataset contains 18082 gene expression measurements on 30365 cells. We remove genes that are expressed in less than 500 cells, normalize the counts per cell to 10000, and log-transform the data. We retain 8000 highly variable genes based on the dispersion score. We let $G$ to be the resulting gene sets.

##### 1.2 Marker Panel Generation for Label-free Methods

Given a predefined marker panel size $k$, we run geneBasis as described in the main paper to obtain the marker panel. PERSIST allows direct specification of $k$ genes to maximally predict the remaining gene expression profile, so we use these $k$ genes as the output panel. For SCMER, we apply a binary search on its regularization parameter until the marker panel size reaches $k$. DUBStepR, however, does not permit direct specification of panel size, so we select the first $k$ genes from its feature set. Additionally, DUBStepR outputs an optimal feature set based on a proxy metric for cell type separation. If the size of the optimal feature set exceeds $k$, we prioritize selecting the first $k$ genes from this set.

#### 2 Specificity Score

We introduce an entropy-based cell-type specificity score. Given gene $g$, let $x_g^{(i)}$ be the expression of gene $g$ in cell $i$, and let $E_{g,k} = \frac{\sum_{i \in S_k} x_g^{(i)}}{|S_k|}$ be the mean expression of gene $g$ in cell type $k$ where $S_k = \{i : Y_i = k\}$. We further transform $E_g = [E_{g,1}, \ldots, E_{g,K}]$ to $q_g$ where $q_{g,k} = \frac{E_{g,k}}{\sum_{l=1}^{K} E_{g,l}}$. The entropy-based specificity score is then computed as

$$\text{Ent-Spec}(g) = -\sum_{l=1}^{K} q_{g,l} \log q_{g,l}$$

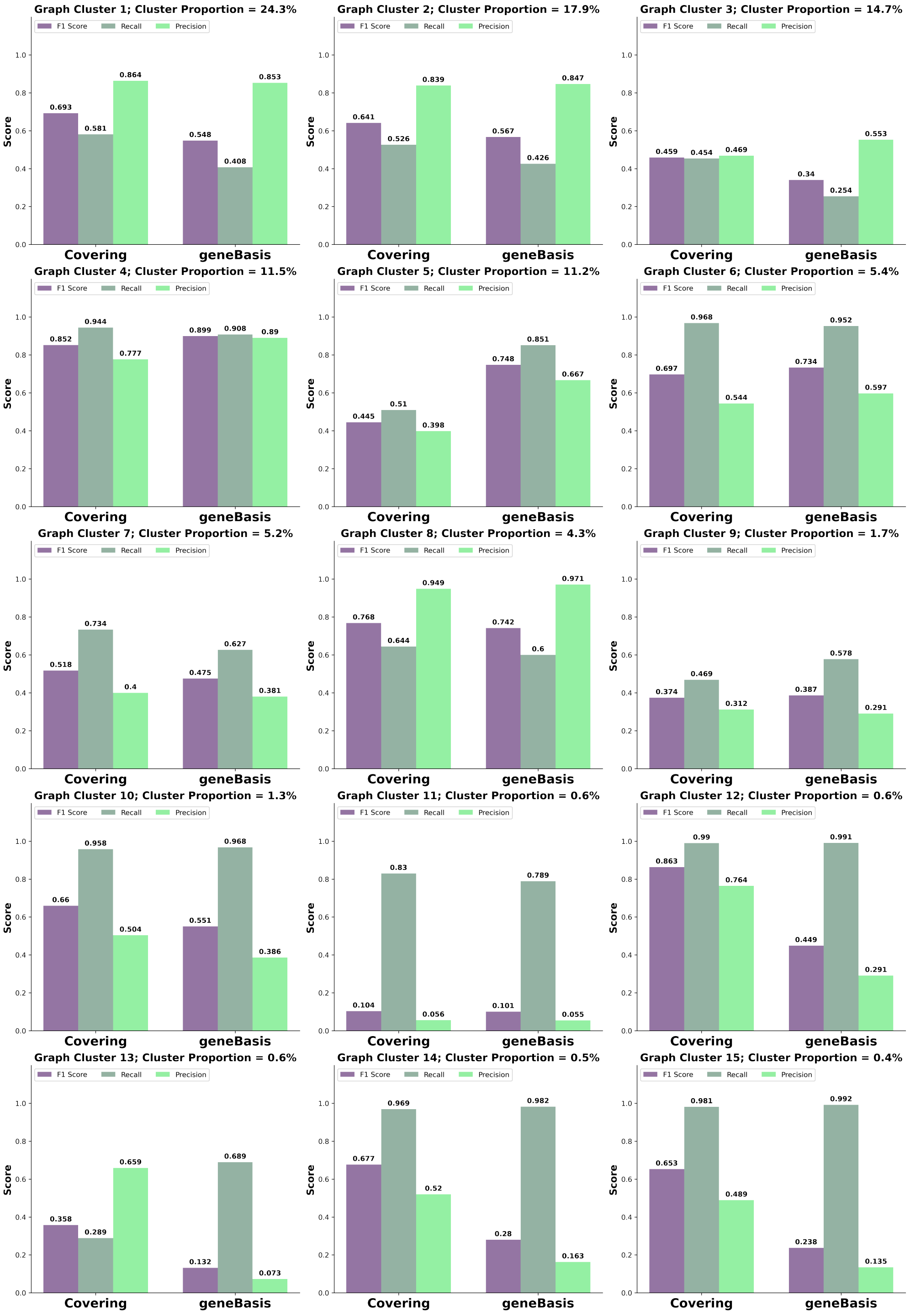


**Fig. 6. Comparison of geneCover and geneBasis in resolving spatial organization in the mouse brain:** We match each cluster from the 10x Genomics (Supplementary Figure 5) with the Leiden clusters learned from the 200 geneCover and geneBasis markers using the procedure described in Supplementary Section 3. The average F1, recall, and precision scores between the 10x Genomics clusters and their best matches over five random seeds of the Leiden algorithm (Supplementary Figure 2A, 2B) are reported.

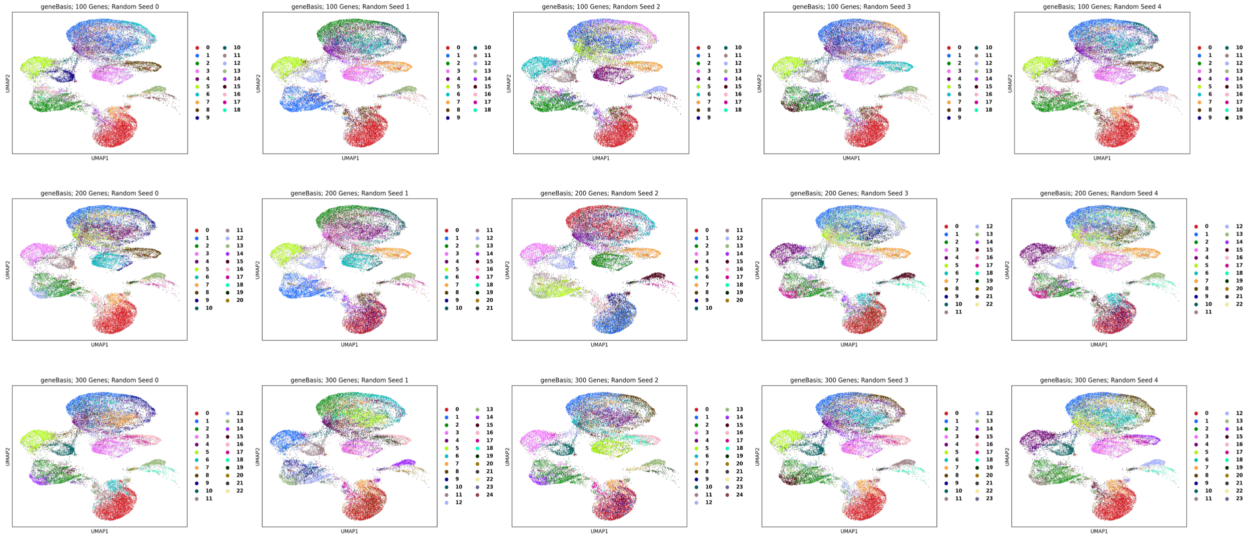


**Fig. 7. Leiden Clusters from GeneBasis Marker Panels with Increasing Sizes Over Multiple Random Seeds:** Leiden clustering is performed on the 50 principal components of the gene expression profile restricted to the geneBasis markers.

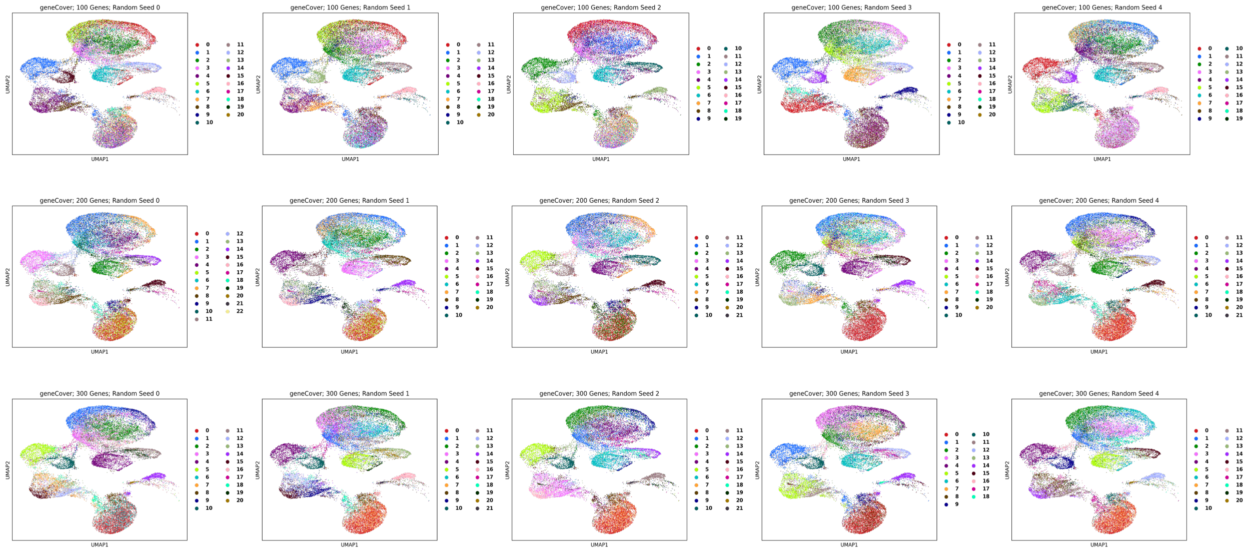


**Fig. 8. Leiden Clusters from GeneCover Marker Panels with Increasing Sizes Over Multiple Random Seeds:** Leiden clustering is performed on the 50 principal components of the gene expression profile restricted to the geneCover markers.

Low score means the marker is specific to some cell type. To evaluate the overall specifity of a marker gene panel, we compute the average of Ent-Spec($g$) across all gene $g$ in the marker panel.

#### 3 Matching Cell Types via F1 Score

Let $Y^* \in \{1, \dots, K^*\}^N$ and $Y \in \{1, \dots, K\}^N$ be reference and target cluster vectors, where $K$ is the number of distinct clusters in $Y$, $K^*$ is the number of distinct clusters in $Y^*$, and $N$ is the total number of cells or spots. For $k^* = 1, \dots, K^*$ (resp. $k = 1, \dots, K$), let $N^*(k^*)$ (resp. $N(k)$) denote the number of cells in $Y^*$ (resp. $Y$) with label $k^*$ (resp. $K$). Let $n(k^*, k)$ be the number of cells with label $k^*$ in $Y^*$ and $k$ in $Y$.

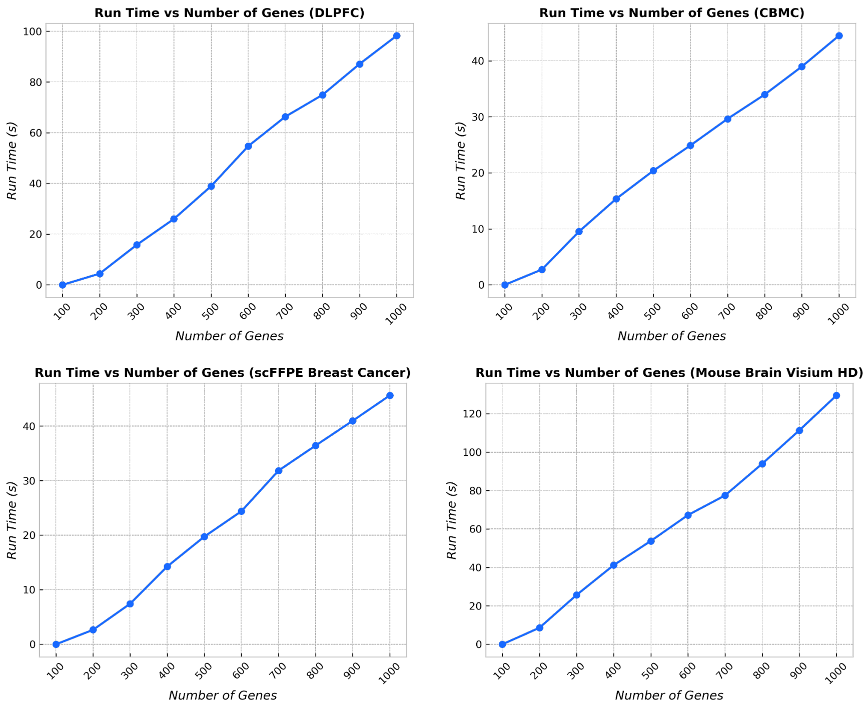


**Fig. 9. Run Time of Iterative GeneCover vs the Gene Panel Size:** The iterative geneCover algorithm is applied to four datasets with $m = 3$ and $\{k_t\}_{1:T} = [100] * T$ for $T = 1, 2, \dots, 10$.

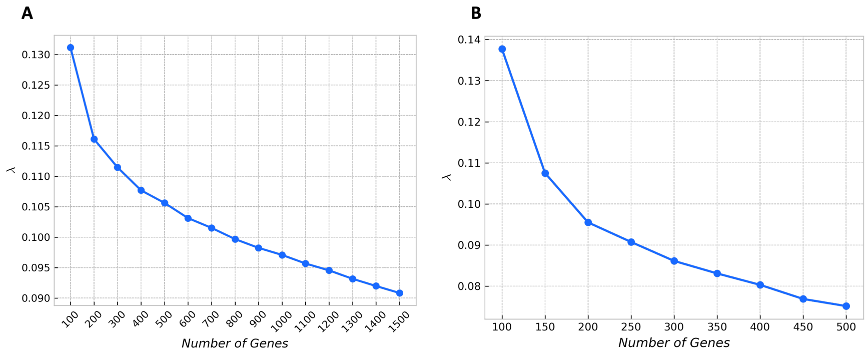


**Fig. 10. λ after Binary Search vs. Gene Panel Size:** For each panel size on the x-axis, the GeneCover algorithm is applied to the dataset using a single run with $m = 3$. Beyond the x-axis limits shown, GeneCover is unable to return the optimal solution within 10 minutes. (A) DLPFC dataset. (B) CBMC dataset.

The F1 score between clusters $k^*$ in $Y^*$ and $k$ in $Y$ being defined by

$$\text{F1}_{Y^*,Y}(k^*, k) = \frac{2n(k, k^*)}{N^*(k^*) + N(k)},$$

we define the best match to cluster $k^*$ in $Y^*$ as the cluster $l$ in $Y$ such that

$$l = \underset{k \in \{1,\dots,K\}}{\arg\max}\ \text{F1}_{Y^*,Y}(k^*, k).$$

We will also compute the precision and recall associated to matching $k$ to $k^*$ as

$$\text{Precision}(k^*, k) = n(k^*, k)/N(k),\ \text{ and Recall}(k^*, k) = n(k^*, k)/N^*(k^*).$$

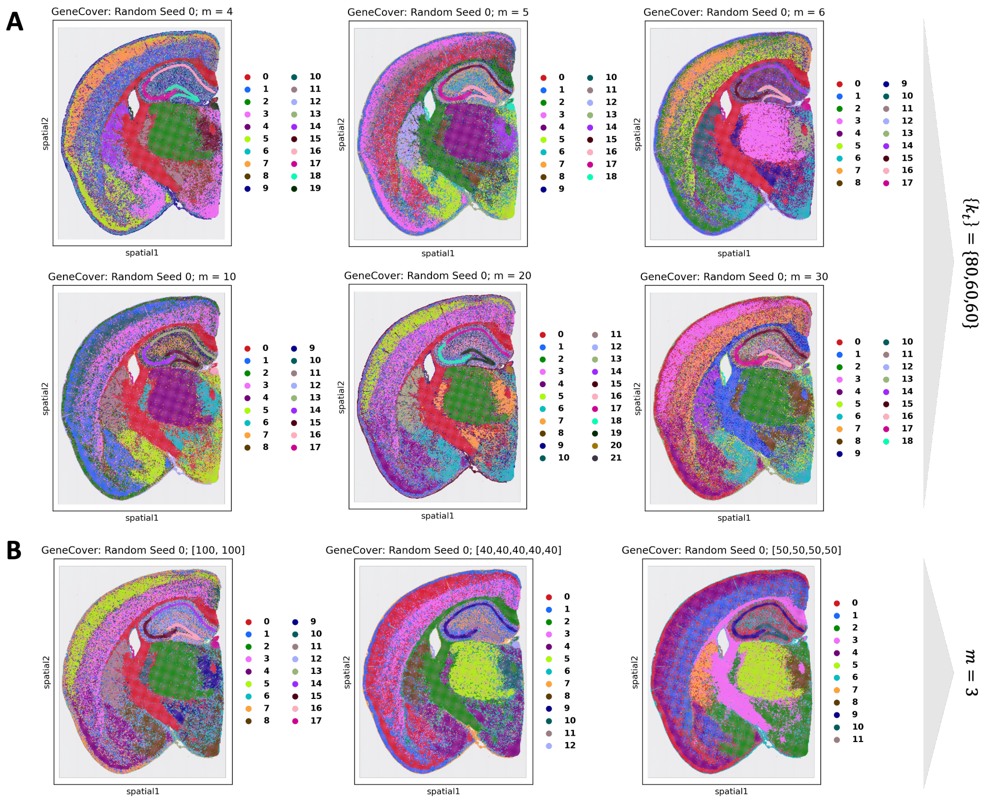


**Fig. 11. Impact of GeneCover Hyperparameter Selection on Clustering Performance:** (A) Leiden clustering results using the GeneCover marker panel generated with $\{k_t\} = [80, 60, 60]$ and increasing $m$, with a fixed clustering resolution of 1.5. (B) Leiden clustering results for a size-200 GeneCover marker panel generated with $m = 3$ and varying incremental sequences, with a fixed clustering resolution of 1.0.

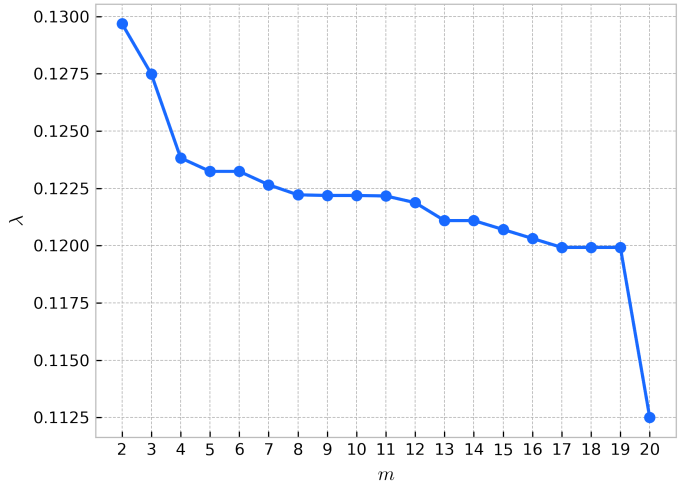


**Fig. 12. $\lambda$ after Binary Search vs $m$ in the Mouse Brain Visium HD Dataset with $\{k_t\}_{1:T} = \{80\}$**

In our analysis, to robustly test the performance of each method, we will obtain multiple $Y$'s using different random seeds of the clustering algorithm. Denote the $Y^{(s)}$ denote the cluster vector from the $s^{\text{th}}$

random seed. For each random seed, suppose we match the cluster $k^*$ in reference $Y^*$ with cluster $l^{(s)}$ in $Y^{(s)}$. Then we will measure the capability of an unsupervised marker gene selection algorithm in recovering reference cluster $k^*$ via the average F1 score between $k^*$ in $Y^*$ and its best match in $Y^s$ over multiple random seeds $s$. In particular, if there are $S$ random seeds in total, then the final metric will be

$$Q_{\mathrm{F1}} = \frac{1}{S}\sum_{s=1}^{S} \mathrm{F1}_{Y^*,Y^{(s)}}(k^*, l^{(s)})$$

We also report the average recall and precision between the reference cluster and its best match over random seeds, denoted $Q_{\mathrm{Recall}}$ and $Q_{\mathrm{Precision}}$.

#### 4 Marker Gene Selection Across Samples

Given a collection of samples $B$, a subset of genes $G \subseteq [\![d]\!]$, and a correlation threshold $\lambda$, we define the neighborhood of gene $j \in G$ within sample $b \in B$ as $M_{G,j}^{\lambda,b} = \{j' \in G : \rho^b(j,j') \geq \lambda\}$, where $\rho^b$ is the Spearman's correlation matrix computed on the rank representation of gene expressions within sample $b$. This correlation structure is encoded in a binary adjacency matrix $A_G^{\lambda,b} \in \{0,1\}^{|G|\times|G|}$ such that $A_G^{\lambda,b}(j,j') = 1$ if $j' \in M_{G,j}^{\lambda,b}$ and $A_G^{\lambda,b}(j,j') = 0$ otherwise. Let $w_g$ reflect the general cost of including gene $g$ to the marker panel, we define the generalized integer programming formulation by:

$$\begin{aligned} &\min_{u} \boldsymbol{w_G} \cdot \boldsymbol{u} \\ &\text{s.t.} \sum_{k\in G} A_G^{\lambda,b}(k,l)u_k \geq 1, \quad \forall l \in G, \forall b \in B \end{aligned} \tag{1}$$

where the objective minimizes the weighted selection of marker genes while ensuring that each gene $l$ in the transcriptome is covered in every sample. The geneCover algorithm and the iterative approach under the new general framework is given below in Algorithm 1.

**Algorithm 1** GeneCover

**procedure** GENERALIZED-GENECOVER($k, \{\boldsymbol{\rho^b}\}_{\boldsymbol{b\in B}}, \boldsymbol{w}, G, m$)
  $J^* = \emptyset$
  $\lambda_{\min} = 0$, $\lambda_{\max} = 1$
  **while** $|J^*| \neq k$ **do**
    $\lambda = \frac{1}{2}(\lambda_{\min} + \lambda_{\max})$
    $A_G^{\lambda,b} = \mathbf{1}_{\boldsymbol{\rho_G^b}>\lambda}, \quad \forall b \in B$ ▷ Binarize $\boldsymbol{\rho_G^b}$ based on threshold $\lambda$
    u = MinimalSetCover($\boldsymbol{w_G}, \{A_G^{\lambda,b}\}_{b\in B}$) ▷ Solve Formulation (1) with weights $\boldsymbol{w_G}$ and matrices $\{A_G^{\lambda,b}\}_{b\in B}$
    $J^* = \{j \in G : u_j = 1, |M_{G,j}^{\lambda,b}| \geq m, \forall b \in B\}$
    **if** $|J^*| < k$ **then**
      $\lambda_{\max} = \lambda$
    **else**
      $\lambda_{\min} = \lambda$
  **Return** $J^*$

**procedure** GENERALIZED-ITERATIVE-GENECOVER($\{k_t\}_{1:T}, \{\boldsymbol{\rho^b}\}_{\boldsymbol{b\in B}}, \boldsymbol{w}, G, m$)
  **for** t = 1: T **do**
    $J_G^* =$ GENERALIZED-GENECOVER($k_t, \{\boldsymbol{\rho^b}\}_{\boldsymbol{b\in B}}, \boldsymbol{w}, G, m$)
    $J^* = J^* \cup J_G^*$
    $G = G \setminus J_G^*$
  **Return** $J^*$
